# Long-term stability of single neuron activity in the motor system

**DOI:** 10.1101/2021.10.27.465945

**Authors:** Kristopher T. Jensen, Naama Kadmon Harpaz, Ashesh K. Dhawale, Steffen B. E. Wolff, Bence P. Ölveczky

**Affiliations:** Department of Organismic and Evolutionary Biology and Center for Brain Science, Harvard University; Computational and Biological Learning Lab, Department of Engineering, University of Cambridge; Centre for Neuroscience, Indian Institute of Science, Bangalore, India; Department of Pharmacology, University of Maryland School of Medicine, Baltimore MD 21201, USA

## Abstract

How an established behavior is retained and stably produced by a nervous system in constant flux remains a mystery. One possible solution is to fix the activity patterns of single neurons in the relevant circuits. Alternatively, activity in single cells could drift over time provided that the population dynamics are constrained to produce stable behavior. To arbitrate between these possibilities, we recorded single unit activity in motor cortex and striatum continuously for several weeks as rats performed stereotyped motor behaviors – both learned and innate. We found long-term stability in single neuron activity patterns across both brain regions. A small amount of drift in neural activity, observed over weeks of recording, could be explained by concomitant changes in task-irrelevant behavioral output. These results suggest that stereotyped behaviors are generated by stable single neuron activity patterns.

## Introduction

### Learning and memory in dynamic motor circuits

When we wake up in the morning, we usually brush our teeth. Some of us then cycle to work, where we log on to the computer by typing our password. After work, we might go for a game of tennis, gracefully hitting the serve in one fluid motion. These motor skills, and many others, are acquired through repeated practice and stored in the motor circuits of the brain, where they are stably maintained and can be reliably executed even after months of no intervening practice^1–3^. The neural circuits underlying such motor skills have been the subject of extensive study^4–8^, yet little is known about how these skills persist over time. Given the stability of the behaviors themselves^9^, a possible solution is to dedicate a neural circuit to a given skill or behavior, then leave it untouched. However, cortical areas undergo continual synaptic turnover even in adult animals^10–13^ and have been shown to change their activity patterns over time, both in the presence and absence of explicit learning^14–18^. While neural circuits in constant flux may facilitate learning of new behaviors and reflect the continual acquisition of new memories and associations^19^, it seems antithetical to the stable storage of previously acquired behaviors.

### Competing theories and predictions

Two main theories have been put forth to explain the apparent paradox of stable memories in plastic circuits. In the commonly held view that motor control is governed by low-dimensional dynamics^20–23^, the paradox can be resolved by having a ‘degenerate subspace’, in which neural activity can change without affecting behavior^24^ or task performance^25^. While this would do away with the requirement for stable activity at the level of single neurons (Figure 1A)^14^, it requires any drift in population activity to occur exclusively in the degenerate subspace. Whether and how biological circuits can ensure this without continual practice remains largely unknown^19^. Absent complete degeneracy, it has also been suggested that the connections from drifting neural populations to downstream circuits could continually rewire to maintain stable motor output and behavior^26^.

**Figure 1:**
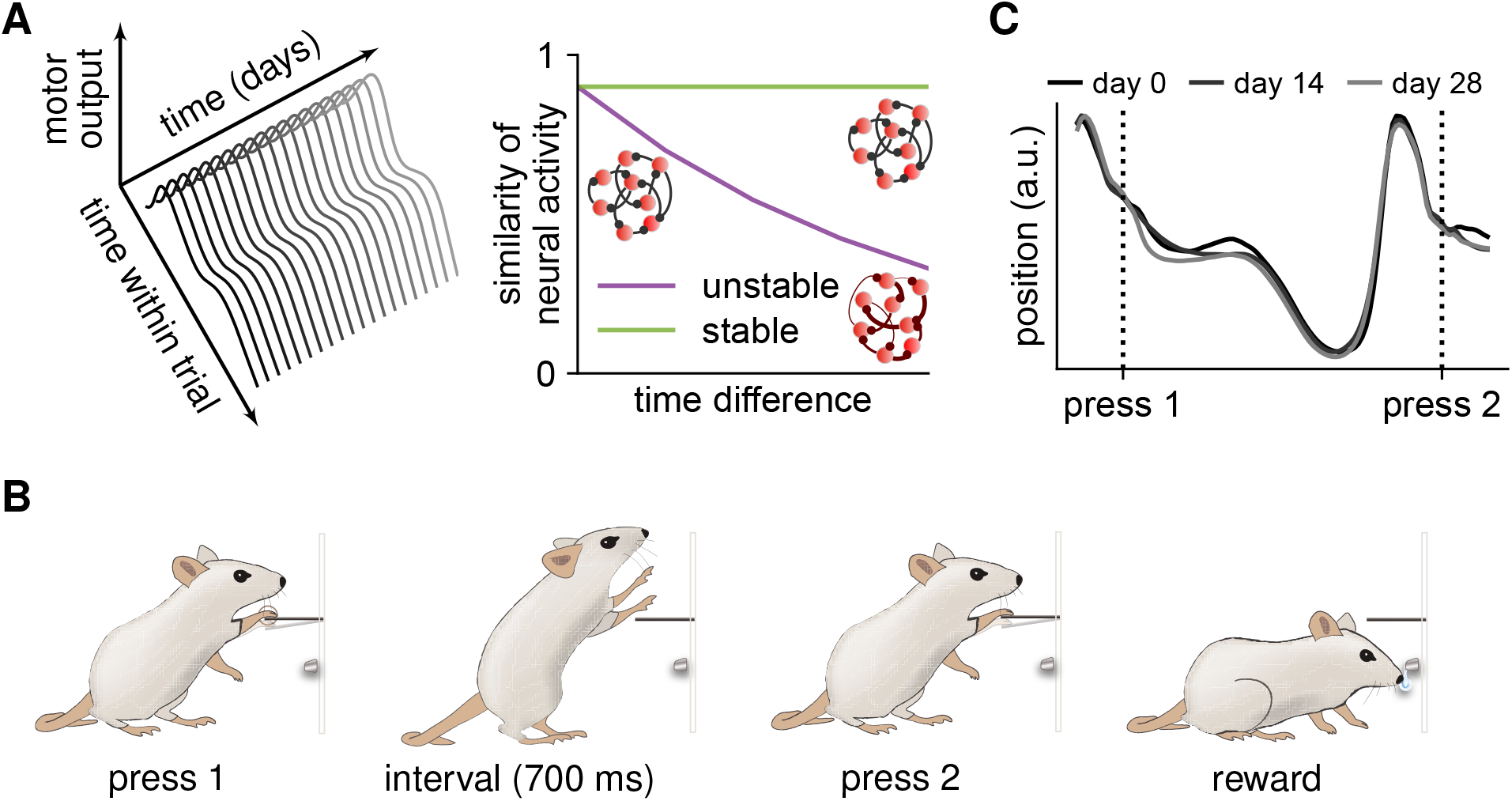
A paradigm for interrogating long-term neural and behavioral stability. **(A)** Schematic illustrating stable and drifting neural activity. For a constant motor output over time (left), the underlying task-related neural activity can either remain stable or change along a behavioral ‘null direction’ (right)^48^. If single neuron activity is stable over time, similarity of the firing patterns associated with two trials of a stable behavior should not depend on the time separating the trials (green). This can be achieved through stable connectivity (RNN insets). Conversely, if the single neuron activity patterns driving the behavior change over time, the similarity of task-associated neural activity will decrease with increasing time difference (purple)^14^. **(B)** Schematic illustration of the task used to train complex stereotyped and stable movement patterns in rats^6^. To receive a reward, rats must press a lever twice separated by an interval of 700 ms. **(C)** Mean task-related forelimb trajectory for an example rat trained on the task in (B) across three different days, each two weeks apart. Y-axis indicates horizontal forelimb position (parallel to the ground).

A different way to maintain stable motor output is by constraining the changes in neural circuits such that they do not affect single neuron activity associated with already established behaviors^25, 27, 28^. In this case, the activity patterns of individual neurons locked to the behavior would remain constant or highly similar over time (Figure 1A)^14^. This solution has been observed in the specialized zebra finch song circuit, where neural activity patterns associated with a stereotyped song remain stable for months^29^. However, zebra finches have a neural circuit dedicated exclusively to learning and generating their one song, with plasticity largely restricted to a ‘critical period’ of development^30^. In contrast, humans and other mammals use the same ‘general’ motor network for a wide range of behaviors – both learned and innate. Whether a similar mechanism could underlie the stability of motor memories in these more generalist circuits has yet to be determined.

It has also been hypothesized that stable circuit function could be associated with stable single neuron activity in some brain regions and constrained population dynamics with drifting single neuron activity in others^19^. In this view, brain regions several synapses removed from the periphery, and with high degrees of redundancy, would be more likely to exhibit representational drift^15, 19, 31^. In contrast, regions closer to the periphery, that serve as information bottlenecks for sensory input and motor output, would maintain more stable representations^14, 31^. However, it is unclear whether the mammalian brain shows such differences in single neuron stability, and, if so, at which stage of the motor hierarchy stable single neuron activity emerges. More generally, it remains an open question whether single unit neural activity patterns in the mammalian motor system are stable over time^24, 32–34^.

### Experimental challenges

Arbitrating between the hypotheses outlined above has been attempted by recording neural activity over time during the performance of well-specified behaviors, either by means of electrophysiology^24, 32, 34–37^ or calcium imaging^15, 29, 38^. These studies have come to discrepant conclusions, with some suggesting stable single unit acitivity^29, 32–34, 37^, and others reporting changing activity for fixed behaviors^24, 35, 38^. It remains unclear whether these discrepancies reflect technical differences in recordings and analyses, or whether they reflect biological differences between behaviors, animals, or circuits as suggested above. Importantly, putative drift in neural activity could be caused by factors not directly related to the mapping between neural activity and motor output. These include unstable environmental conditions or fluctuations in the animal’s internal state, including attention, satiety, and motivation^39–41^. Notably, many of these processes, driven by constrained or cyclic fluctuations in hormones or neuromodulators^40, 42^, drift around a mean. They can therefore be distinguished from drift in neural circuits by recording for durations longer than the autocorrelation time of the various uncontrolled, or ‘latent’, processes.

However, high-quality long-term recordings of the same neurons can be technically challenging. In lieu of this, a recent approach has considered the stability of low-dimensional latent neural dynamics over extended time periods. This was done for motor cortex by applying linear dimensionality reduction to recordings from each experimental session followed by alignment of the resultant low-dimensional dynamics^43^. While this work suggests that latent motor cortical dynamics underlying stable motor behaviors are stable over time, it does not address the source of this stability. In particular, it remains unclear whether such long-term stable latent dynamics result from drifting single-unit activity within a degenerate subspace that produces the same latent trajectories, or whether it is a consequence of neural activity patterns that are stable at the level of single units.

In this work, we first use a recurrent neural network to demonstrate how long-term single-unit recordings during a stably executed behavior can distinguish between the two main models of how stable behaviors are maintained. We then go on to perform such recordings in rats producing stable behaviors, considering two central nodes of the motor system: motor cortex (MC) and dorsolateral striatum (DLS)^44^. Importantly, both MC and DLS have orders of magnitude more neurons than the lower-level control bottlenecks they project to^45–47^, and they therefore exhibit substantial degeneracy with respect to motor output^47, 48^. To probe the degree to which our findings generalize across different classes of behaviors relying on different control circuits, we examine both learned (Figure 1B) and innate behaviors. To minimize sources of neural variability not directly related to behavioral control, we performed our experiments in a highly stable and controlled environment. Additionally, we recorded the animals’ behavior at high spatiotemporal resolution to account for any changes in task-irrelevant movements (Figure 1C)^32, 49^. Our combined neural and behavioral recordings revealed that neural circuit dynamics are highly stable at the level of single neurons. The small amount of drift in task-related neural activity could be accounted for by a concomitant slow drift in the behavior. These results suggest that stable behaviors are stored and generated by stable single-unit activity in the motor circuits that drive the learned behavior, and that the neural correlates of behavior are also stable in an innate behavior, which does not directly depend on the motor circuits we record from.

## Results

### Network models of stable and unstable motor circuits

When analyzing the stability of task-associated neural activity, it is important to consider stability not only at the population level (e.g. in the form of stable latent dynamics), but also at the level of single task-associated neurons. We start by simulating a degenerate artificial control network to highlight this distinction and motivate the use of longitudinal single-unit recordings to address the neural mechanisms of long-term behavioral stability. Our simulations show how ‘stable’ and ‘drifting’ single neuron activity can both drive stable latent dynamics and behavior, providing validation and motivation for the experimental strategy and analyses considered in this work^50^. Our simulated neural circuit was a recurrent neural network (RNN) producing a stereotyped output, akin to those previously used to model pattern generator functions^51–54^ (Methods). We trained the network using gradient descent^55^ to generate five smooth target output trajectories (Figure 2A; Methods). After training, we simulated the noisy dynamics for 100 trials (Methods) and generated spikes from a Poisson observation model to constitute a simulated experimental ‘session’ (Figure 2B, 2C).

**Figure 2:**
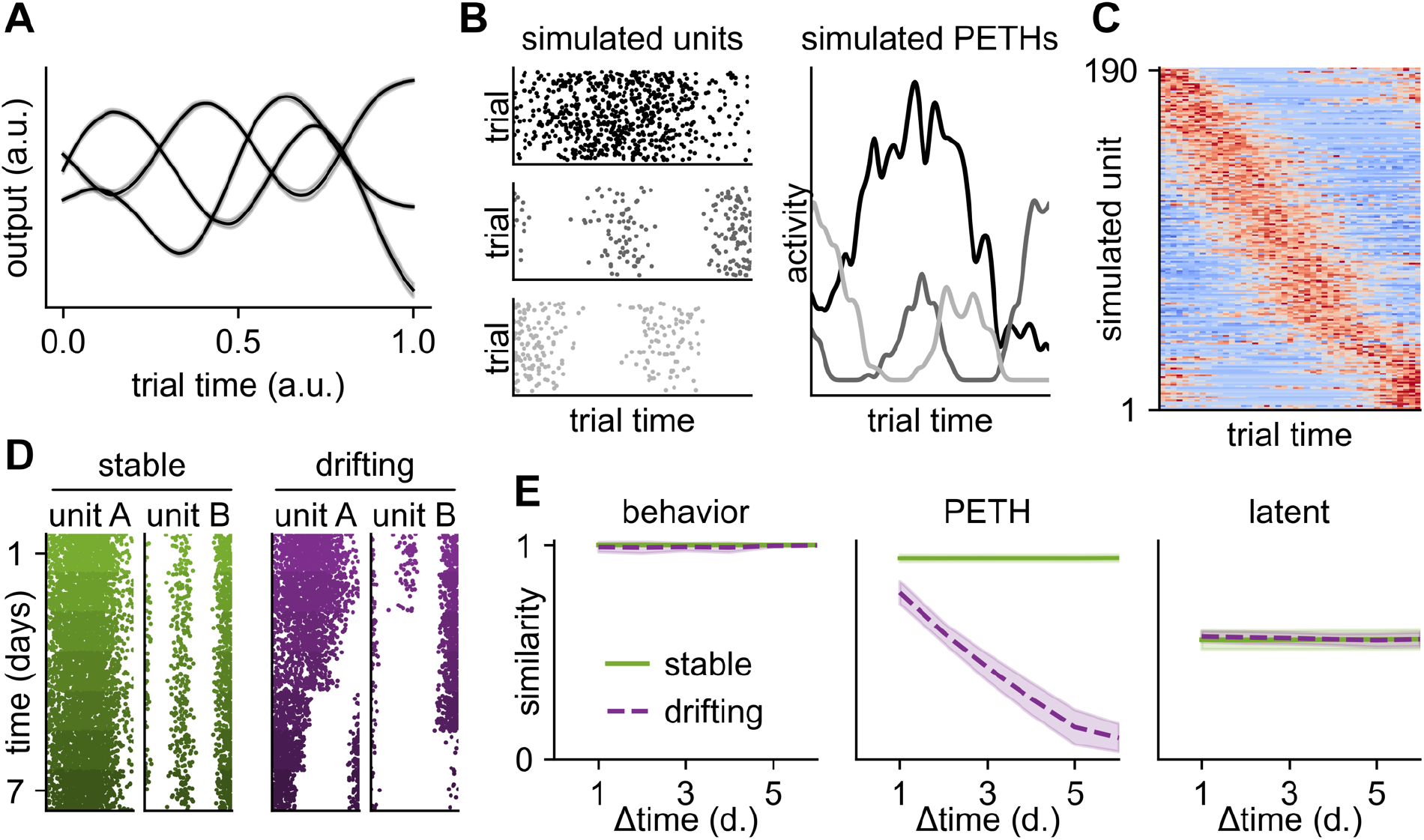
Analyzing neural stability in a recurrent network model. **(A)** Example RNN outputs after training (mean across 100 simulated trials). **(B)** Activity of three example recurrent neurons after training (Methods), visualized as raster plots of spike times (left) or peri-event time histograms (PETHs; right) across 100 simulated trials. **(C)** PETHs for all units firing at least 100 spikes, sorted according to the PETH peak from a set of held-out trials and plotted as a heatmap with color indicating spike count from low (blue) to high (red). **(D)** Example raster plots as in (B) across 7 different sessions (y-axis) for a network exhibiting either stability (left) or drifting neural activity (right). **(E)** Quantification of the similarity in the space of network output (left), PETHs (middle), and aligned latent trajectories (right) as a function of time difference (change in y value from (D)) for the stable RNN (green) and the drifting RNN (purple). Lines and shadings indicate mean and standard deviation across 10 networks.

Due to the degeneracy of the circuit (250 neurons with 60,000 parameters controlling a 5-dimensional time-varying output), multiple distinct networks with different single-unit activity patterns can achieve the same target output. This allowed us to compare network dynamics of RNNs producing the same output with either identical or differing connectivity across separate simulated ‘sessions’. To intuit how activity patterns change over time in an unstable network, we performed a linear interpolation between the parameters of two independently trained RNNs and finetuned the networks to ensure robust performance (Methods). This yielded 7 RNNs with progressively more dissimilar connectivity, yet which all produced the same output – a phenomenological model of neural drift, where the position of a network within this interpolation series is used as a proxy for time (Extended Data Figure 1). We proceeded to investigate the degree to which single-unit activity changed as a function of this measure of time. When inspecting the activity of individual units in the RNNs, we found that their firing patterns tended to change from session to session, with sessions close in time generally exhibiting more similar firing patterns than distant sessions (Figure 2D). To quantify this, we computed the correlation between single-unit PETHs for all pairs of sessions, averaged across units. This PETH correlation systematically decreased as a function of time difference between sessions, despite a stable network output (Figure 2E). In contrast, a negative control where network parameters fluctuated around a single local minimum yielded single-unit neural activity that was highly stable over time (Figure 2E; Methods).

Finally, we considered how such single-unit analyses differ from approaches that consider the stability of low-dimensional latent dynamics^43^. Following Gallego et al^43^, we computed the principal components of neural activity for each session and aligned the resulting latent dynamics by applying canonical correlation analysis (CCA) to each pair of simulated sessions (Methods). We then computed the neural similarity as a function of time difference, measuring similarity as the correlation between aligned latent trajectories. As expected for networks with constant output, the latent dynamics did not become more dissimilar over time for either of the networks with constant or drifting single unit activity (Figure 2E). These observations reflect the fact that the drifting network retained stable population statistics due to the conserved nature of the task, despite the changing activity profiles of individual units^56^. Our artificial model system thus highlights the importance of long-term recordings of single units to complement studies of latent space stability when investigating how stable behaviors are generated. In particular, we illustrate how stability at the level of single units implies stability at the level of latent trajectories, while the stable latent dynamics reported in previous work^43^ can be driven by either stable or drifting single-unit activity patterns.

### Long-term recordings of neural activity and kinematics during a learned motor task

To investigate the stability of biological motor circuits experimentally, we trained rats (n=6) to perform a timed lever-pressing task in which they received a water reward for pressing a lever twice with an inter-press interval of 700 ms. Rats learned to solve the task by developing complex stereotyped movement patterns (Figure 3A, Extended Data Figure 2A)^6, 57^. Since the task is kinematically unconstrained (meaning it has many ‘motor solutions’) and acquired through trial-and-error, each animal converged on its own idiosyncratic solution (Figure 3B). However, once acquired, the individually distinct behaviors persisted over long periods of time (Figure 3A).

**Figure 3:**
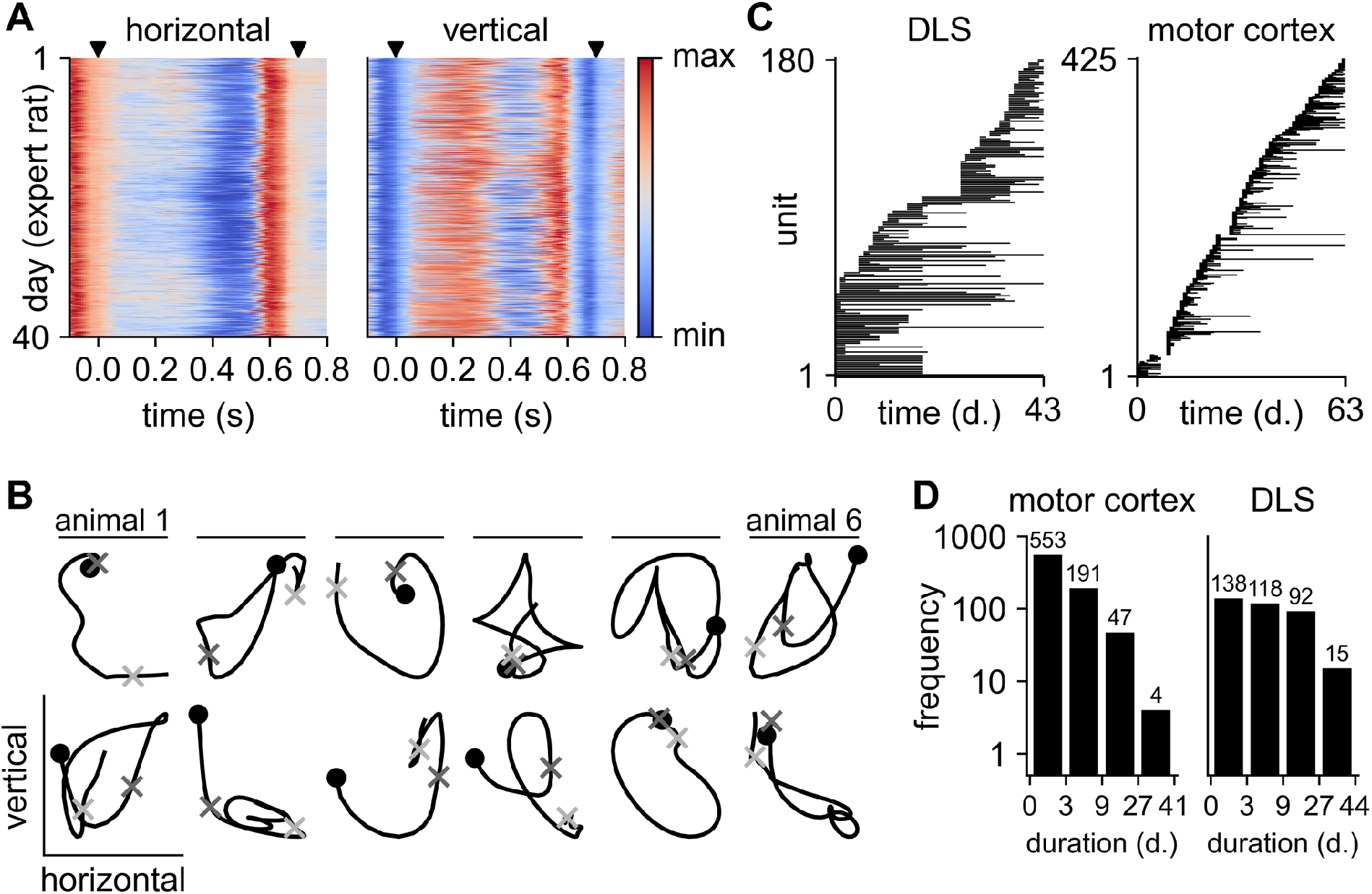
Experimental recordings of behavior and neural activity. **(A)** Right forelimb trajectories in the horizontal and vertical directions (c.f. Figure 1B) for an example expert rat (see Extended Data Figure 2A for data from the other 5 animals). Color indicates forelimb position. Kinematics were linearly time-warped to align the two lever-presses for all analyses (Methods; warping coefficient = 1.00 ± 0.07), and black triangles indicate the times of the lever presses. The rat uses the same motor sequence to solve the task over many days with only minor variations. **(B)** Mean trajectories across all trials of the left (top row; left side view) and right (bottom row; right side view) forelimbs for each rat (columns), illustrating the idiosyncratic movement patterns learned by different animals to solve the task. Circles indicate movement initiation; dark and light grey crosses indicate the times of the 1^st^ and 2^nd^ lever press respectively. **(C)** Time of recording for each unit for two example rats recording from DLS (left) and MC (right). Units are sorted according to the time of first recording. **(D)** Distribution of recording times pooled across units from all animals recording from DLS (left) or MC (right). Numbers above bars indicate the number of neurons in each bin. Note that the data used in this study has previously been analyzed by Dhawale et al.^57, 59^.

To reduce day-to-day fluctuations in environmental conditions that could confound our assessment of neural stability over time, animals were trained in a fully automated home-cage training system with a highly regimented training protocol in a very stable and well-controlled environment^58^. After reaching expert performance, animals were implanted with tetrode drives for neural recordings^59^ targeting motor cortex (MC) or dorsolateral striatum (DLS)^44^ (Methods). While the stability of single units in cortical regions has previously been addressed with inconsistent findings^14, 24, 32, 33, 59^, studies of neural stability in sub-cortical regions, and specifically the striatum, are scarce^60, 61^. DLS is, in this case, particularly relevant as it is essential for the acquisition and control of the motor skills we train^57^.

Three animals were implanted in Layer 5 of MC, and three animals in DLS. Following implantation and recovery, animals were returned to their home-cage training boxes and resumed the task. Neural activity was then recorded continuously over the course of the experiment^59^. Importantly, our semi-automated and previously benchmarked spike-sorting routine^59^ allowed us to track the activity of the same neurons over days to weeks in both DLS and MC (Figure 3C, 3D). The task-relevant movements of all animals were tracked using high-resolution behavioral recordings^62, 63^, and both behavior (kinematic features) and neural activity were aligned to the two lever-presses to account for minor variations in the inter-press interval (Methods).

### Single neurons in MC and DLS have stable activity patterns

The combination of controlled and regimented experimental conditions, stable behavior, and continuous neural recordings provides a unique setting for quantifying the stability of an adaptable circuit driving a complex learned motor behavior^14^. Importantly, this experimental setup mirrors the scenario considered in our RNN model (Figure 2) and thus facilitates analyses of neural stability at the level of single neurons. We first considered the PETHs of all units combined across all trials and found that units in both MC and DLS fired preferentially during particular phases of the learned behavior (Figure 4A)^59^. Importantly, we found that the behaviorally locked activity profiles of individual units appeared highly stable over long periods of time (Figure 4B), reminiscent of the ‘stable’ RNN model (Figure 2D).

**Figure 4:**
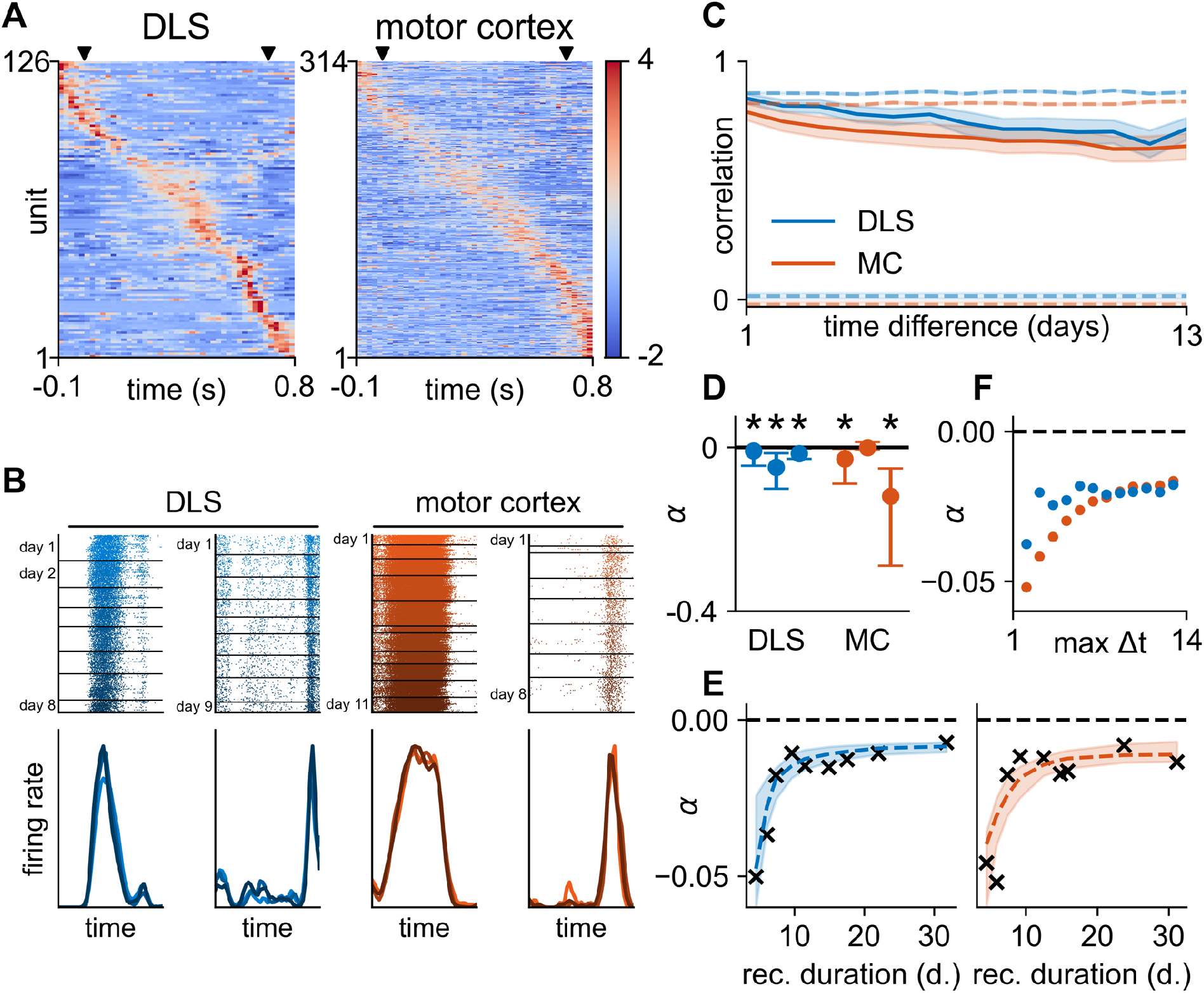
Single-unit activity is stable over time in DLS and MC. **(A)** z-scored PETHs across trials and sessions for all units firing at least 100 spikes during the lever-pressing task for two example rats. Units were sorted according to the activity peak from a set of held-out trials. X-axis indicates time-within-trial relative to the first lever press, spike times were linearly time-warped to align the two lever presses (Methods; see Extended Data Figure 7 for results without time-warping), and black triangles indicate the times of the presses. **(B; top)** Raster plots for two example units in DLS (left) and MC (right) illustrating firing patterns that are time-locked to the behavior over days to weeks. Horizontal lines indicate the beginning of a new day, and color indicates the progression of time from day 1 (light) to the last day of recording (dark). **(B; bottom)** Normalized PETHs for the four example units computed on three different days (early, middle, late) with corresponding colors in the raster. Our quantification of neural similarity is based on the correlations between such PETHs. **(C)** Mean value of the correlation between PETHs calculated on separate days, averaged over all units recorded for at least 14 days from MC (red) or DLS (red) and plotted as a function of time between days (n = 38 neurons for DLS; n = 27 neurons for MC; see Extended Data Figure 3A for data from individual neurons and Extended Data Figure 4A for different recording thresholds). Shaded regions indicate standard error across units. Colored dashed lines indicate the similarity between non-identical neurons (lower) and in a resampled dataset with neural activity drawn from a stationary distribution (upper; Methods). **(D)** Quartiles of the distribution of stability indices for each animal. Horizontal dashed line indicates *α* = 0. Asterisks indicate p < 0.05 for the median stability index being smaller than zero (permutation test; Methods; n = [88, 22, 13] neurons for DLS; [122, 6, 7] for MC). **(E)** Rolling median of the stability index (crosses) for units recorded for different total durations (x-axis). Bins are overlapping with each neuron occurring in two bins (see Extended Data Figure 6C for the non-binned data). Dashed lines indicate exponential model fits to the non-binned data, and shadings indicate interquartile intervals from bootstrapping the units included in the model fits (Methods). **(F)** Stability indices of models fitted to increasing subsets of the data from (C), illustrating how longer recording durations lead to longer time constants. The maximum time difference considered for the model fit is indicated on the x-axis (see Extended Data Figure 8 for the full model fits).

To quantitatively compare neural activity profiles across days, we constructed PETHs for each neuron by summing the spike counts across all trials on each day and convolving them with a 15 ms Gaussian filter (Figure 4B; Methods). We then computed the Pearson correlation *ρ* between pairs of PETHs constructed from neural activity on different days as a function of the time difference between days (Extended Data Figure 3A). This is similar to our RNN analyses (Figure 2) and previous studies in visual and motor circuits^56, 59^. When considering neurons recorded for at least two weeks, the mean PETH similarity remained high in both DLS and MC (Figure 4C; see Extended Data Figure 4A for other recording thresholds). This is consistent with results from the stable RNN model (Figure 2E), and it suggests that learned motor behaviors are driven by single neuron activity patterns that do not change over the duration of our recordings, despite the life-long structural and functional plasticity in these circuits^11, 12, 17, 64^.

To see how this compares to a hypothetical circuit where population statistics are retained but individual neurons change their firing patterns, we also computed pairwise correlations between non-identical neurons recorded on different days. These correlations were near zero in both MC and DLS, confirming that the high correlation over time for individual units is not due to a particular population structure of neural activity imposed by the task (Figure 4C). These results suggest that single neuron activity associated with the learned motor skill is qualitatively stable over periods of several days and weeks (Figure 4B, 4C). Our findings also suggest that the stable latent dynamics identified in previous work^43^ could be a result of such stable single-unit dynamics (c.f. Figure 2E), which is further supported by the fact that alignment of the neural dynamics using CCA did not increase stability (Extended Data Figure 5A).

In contrast to our RNN model, the experimental data contained neurons that were recorded for different durations (Figure 3D). This introduces additional variability and makes it difficult to assess stability across neurons without either discarding neurons recorded for short durations or losing information about neurons recorded for long durations. To combine information across more neurons, we instead considered the PETH similarity as a function of the time difference between PETHs for each neuron individually. An exponential model of the form *ρ* = *βe^αδt^* was fitted to the Pearson correlation (*ρ*) between PETHs as a function of time difference *δt* for each neuron (Methods; see Extended Data Figure 6A for example fits). *α* = −*τ*^-1^ is denoted as the ‘stability index’, since it corresponds to the negative inverse time constant *τ* in an exponential decay model, and this stability index provides a single parameter summarizing the rate of drift for each neuron.

We then considered the distribution of stability indices across neurons recorded for at least 4 days. In a null-model, where single neuron activity remains constant, the PETH similarity should be independent of the time difference for all units (c.f. Figure 2E). The stability indices should thus be centered around zero with some spread due to trial-to-trial variability, corresponding to an infinitely slow exponential decay. The population-level distributions over *α* were indeed centered near zero (Figure 4D). However, a permutation test across time differences revealed that all DLS recordings and two of the animals with recordings from MC did in fact exhibit slow but significant neural drift (p < 0.05). We saw this also when combining data for all neurons across animals within each experimental group (DLS: α*_median_* = −0.014, τ*_median_* = 71 days, p < 0.001; MC: α*_median_* = −0.027, τ*_median_* = 37 days, p < 0.001; permutation test).

### Short recording durations underestimate the stability of neural activity

Our analyses at the level of single neurons included units recorded for short durations of time. However, as noted in the introduction, recording over such short time spans could underestimate stability in the presence of latent processes that affect neural dynamics. Such processes may vary over timescales of hours or days^65^ but be constrained over longer timescales by homeostatic mechanisms, biological rhythms, or task constraints^40–42^. Even though such bounded physiological fluctuations will manifest as short-term drift in neural firing patterns, their contributions to estimates of neural stability will diminish as neural recording durations exceed the characteristic timescales of the underlying processes.

To better estimate drift over longer timescales, we binned the stability indices of all neurons by their recording duration. This revealed that the apparent stability ranged from *α* ≈ −0.05 for short recording durations to *α* ≈ −0.01 for long recording durations (Figure 4E). To extrapolate to longer recording durations, we fitted an exponential model to the data of the form *α* = −*a* − *be*^-*c t*^ (Figure 4E). The parameter *τ*_∞_ = *a*^-1^ provides an estimate of the asymptotic stability of the population and took values of *τ*_∞_ = 139 days for DLS and *τ*_∞_ = 92 days for motor cortex (interquartile ranges of 115-211 for DLS and 75-161 for MC; bootstrapped model fits; Methods). To confirm that the increase in apparent stability with recording duration was not due to a bias in our data collection, we returned to the average similarity across neurons recorded for at least 14 days (Figure 4C). We subsampled the data from these neurons to different maximum time differences, thus varying the effective recording duration for a fixed set of neurons. We then computed stability indices by fitting our exponential model to the average correlation across neurons as a function of time difference (Methods; Extended Data Figure 8). The stability indices increased with subsampled recording duration in both DLS and MC (Figure 4F), consistent with the results across the whole population of neurons (Figure 4E). These findings suggest the presence of constrained fluctuations in the physiology or behavior of the animals, which affect estimates of neural stability on shorter timescales. Our results therefore motivate long-duration single neuron recordings for estimating long-term neural stability.

Finally, if the observed increase in stability with recording duration is due to latent processes with autocorrelations on the order of days, we would expect the neural similarity to decrease to some saturating baseline value, *γ*. We therefore proceeded to fit a model to the average similarity across neurons as a function of time difference, which assumes a decay to such a baseline (*ρ* = *βe^αδt^* + *γ*; Extended Data Figure 8). This model yielded an asymptotic correlation of *γ* = 0.61 for DLS and *γ* = 0.62 for MC. Such positive saturating values suggest a high degree of neural similarity at long timescales.

In summary, we find that estimates of stability can be biased by short recording durations, possibly due to the presence of physiological or behavioral processes with characteristic timescales on the order of hours and days. When considering longer recording durations, we find stability on timescales upwards of 100 days in a single-timescale exponential model (Figure 4E). When instead considering a three-parameter model that included a fitted baseline value *γ*, we found evidence of a saturating similarity substantially larger than the near-zero correlations expected from a complete shuffling of the neural population (Figure 4C). These findings suggest that motor memories are retained by maintaining stable task-associated activity patterns in single neurons.

### Neural drift is correlated with behavioral changes

In the previous section, we quantified the stability of motor circuits assuming a stable behavior and showed how such estimates can be affected by internal or external processes that affect neural activity. However, any residual drift in neural activity is still likely to be an underestimate of the true stability in the mapping from circuit activity to motor output. Indeed, even a perfectly stable neural system should exhibit drifting neural activity patterns locked to the behavior if the behavior itself is changing^32, 66^. This tends to be the case even after performance saturates and stabilizes in a motor task. In particular, humans and animals alike exhibit small behavioral changes both in terms of trial-to-trial variability^67^ and systematic drifts in the mean behavioral output^66^. If such systematic behavioral drift is present in the motor task we analyze, it could explain some of the short-timescale constrained drift as well as any residual drift in neural activity over longer timescales (Figure 4). This, in turn, would suggest a more stable circuit linking neural activity to behavior than revealed by analyses of neural data alone. Thus, to quantify the degree to which the neural drift we see can be accounted for by accompanying changes in task-related motor output, we proceeded to analyze the kinematics of the timed lever-pressing behavior and how they changed over time.

We first examined whether minor behavioral changes could be observed in the motor output by visualizing the z-scored forelimb velocities; that is, we subtracted the mean velocity across all trials for each time point and normalized by the standard deviation. This discarded the dominant mean component of the motor output and revealed a slow drift in the behavior over periods of days to weeks (Figure 5A, Extended Data Figure 2B). To quantify this drift, we computed the correlation between mean forelimb velocities across trials on a given day as a function of the time separating each pair of days. This confirmed the presence of a small but consistent decrease in the similarity of motor output as a function of time difference (Figure 5B; Extended Data Figure 9). Importantly, such behavioral drift occurred despite stable task performance (Extended Data Figure 9). This is consistent with previous work considering behavioral variability in expert performers as an underlying ‘random walk’ in behavioral space with added motor noise^66^, although our work considers drift over the course of several weeks rather than hours. If the physical environment remains unchanged, any long-term behavioral drift must ultimately arise from changes in neural activity. Additionally, DLS is known to be involved in driving the behavioral output during this lever pressing task^57^. These considerations suggest that the observed drift in neural activity could be in directions of state space that affect motor output and thus reflect these changes in behavior. To investigate this, we followed previous work^15, 32^ and showed that the performance of a decoding model predicting behavior from neural activity and an encoding model predicting neural activity from behavior did not deteriorate over time (Extended Data Figure 5). However, the decoding analysis only considered a small subset of our data, where a large population of neurons was recorded simultaneously (c.f. Figure 3C), and it has previously been shown that encoders and decoders can suffer from omitted variable bias^68, 69^. We therefore proceeded to investigate the relationship between neural and behavioral drift at a single neuron level without relying on such estimates of ‘effective connectivity’.

**Figure 5:**
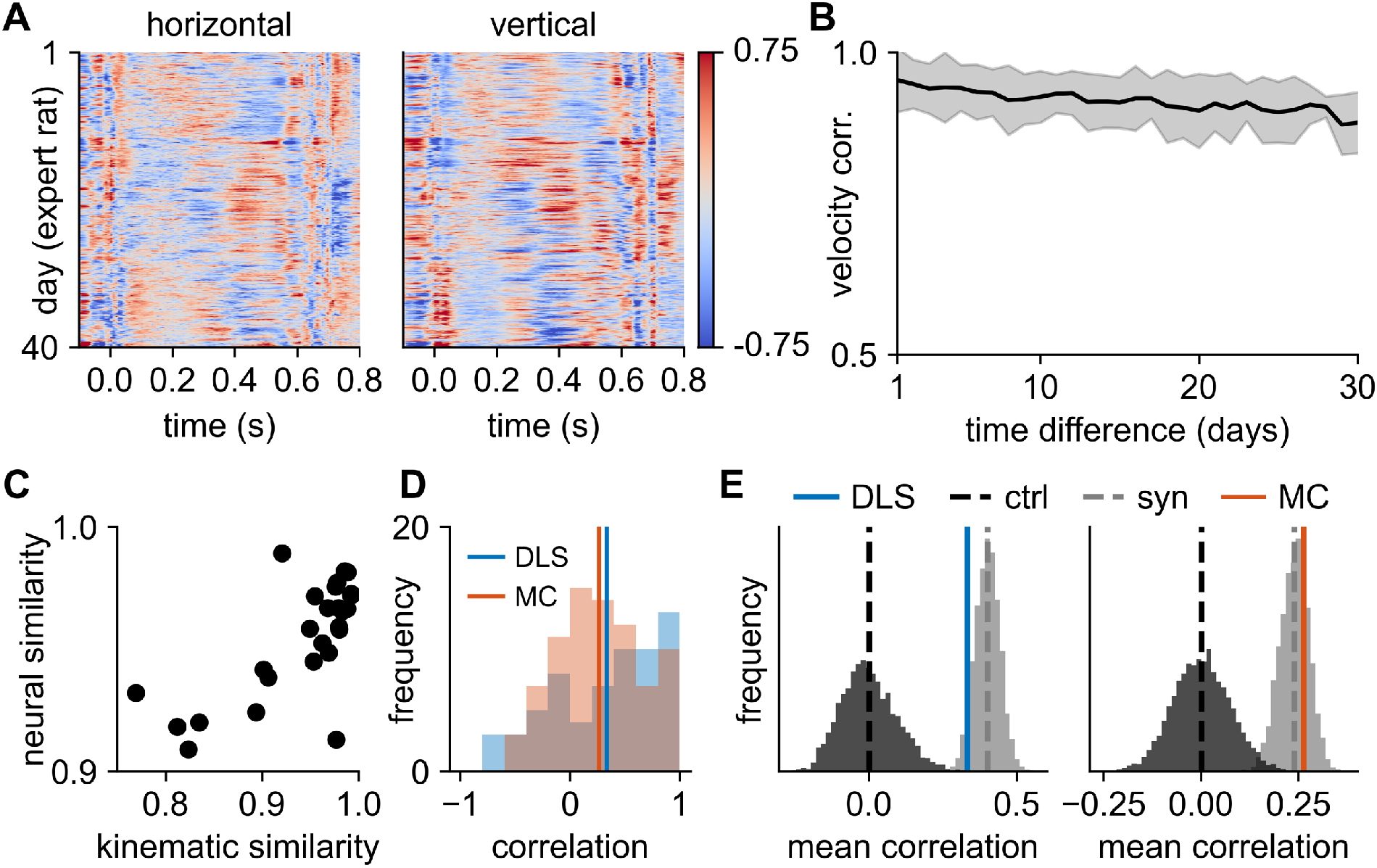
Long-term drift of task specific movement patterns in the lever-pressing task. **(A)** Forelimb velocities for the example animal from Figure 3A, plotted as z-scores with the column-wise mean subtracted (see Extended Data Figure 2B for the other 5 animals). The stereotyped motor sequence masks a behavioral drift across days and weeks. **(B)** Mean and standard deviation of behavioral similarity as a function of time difference, averaged across all pairs of days for the example animal in (A) (see Extended Data Figure 9 for data across all animals). **(C)** Similarity between PETHs on consecutive days plotted against the similarity in kinematic output across the corresponding days for an example unit. Each point corresponds to a single pair of consecutive days. **(D)** Distribution of the correlation between neural similarity and behavioral similarity on consecutive days for neurons recorded in DLS (blue) and MC (red). Vertical lines indicate average correlations. **(E)** Mean correlation between neural and behavioral similarity across neurons from (D), recorded in either DLS (blue; left panel) or MC (red; right panel). Dark grey histograms indicate control distributions constructed by permuting the days in the behavioral data. Light grey histograms indicate the distributions of correlations in synthetic datasets where neural activity is determined entirely by behavior via a GLM.

To do this, we computed both the similarity of neural PETHs and the similarity of forelimb velocity profiles for each pair of consecutive days. We then exploited the fact that the behavioral output changes to different extents on different days (Figure 5A, 5B) and computed the correlation between neural and behavioral drift rates across all consecutive days for each neuron. This correlation should be positive if drift in neural activity is related to drift in motor output (Figure 5C). The mean of the distribution of correlations over all neurons was 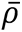 = 0.33 for DLS and 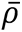 = 0.26 for MC (Figure 5D). These values were significantly larger than null distributions generated by permuting the behavioral data to break any correlations with the neural drift (Figure 5E; p < 0.001; permutation test). This finding confirms that the shorter timescale drift in neural activity is directly related to changes in behavior and suggests that neural drift could be even slower for behaviors with stronger kinematic constraints.

We proceeded to investigate how this experimental correlation compared to a hypothetical system where the drift in neural activity was driven entirely by the drift in behavior, i.e., where there was a stable mapping between single unit activities and behavioral output. To do this, we fitted a linear-nonlinear Poisson GLM^70^ to predict neural activity from behavior using data from a single day of recording for each unit (Methods). This model was used to generate synthetic neural activity on each trial from the recorded behavior, allowing us to compute the correlation between the simulated neural drift and the experimentally observed behavioral drift. Here, we found an average correlation with behavior of 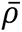 = 0.40 for DLS and 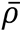 = 0.24 for MC. The correlation values found in the experimental data were substantially more similar to this stable synthetic circuit than to the null distribution with no relation between the drift in neural activity and behavior (Figure 5E; p = 0.06 and p = 0.73 for the synthetic average correlation being smaller than the experimental value).

Taken together, our behavioral analyses confirm that the drift in neural activity is driven, at least in part, by a concomitant task-irrelevant behavioral drift. Additionally, the correlation between the drift in neural activity and behavior is comparable to a synthetic system where the systematic changes in neural activity are exclusively caused by behavioral changes. This suggests that behavioral changes can account for much of the experimentally observed neural drift.

### Neural activity remains stable during an innate behavior

The majority of studies on neural stability have considered behaviors that are either learned or adapted to artificial settings, such as navigating a maze^15^, reaching for points on a screen^24, 32, 34^, controlling a BCI^34, 35, 37^, or singing a song^29, 38^. However, many of the behaviors we express are species-typical, or ‘innate’. For example, sneezing, crying, and shivering require intricate patterns of sequential muscle activity but are not consciously controlled or learned. While we know less about the neural circuits controlling such innate behaviors, we can probe the stability with which they are encoded and compare them to behaviors that explicitly require plasticity. We therefore considered an innate behavior in the rat known as the ‘wet-dog shake’ (WDS), which is characterized by whole-body oscillations^71–74^. Importantly, while we know that MC and DLS are necessary for learning^6^ and executing^57^ the stereotyped motor patterns required for mastering the lever-pressing task, the wet-dog shake is generated by circuits downstream of DLS and MC^75^. If degenerate or redundant circuits exhibit a higher degree of drift for a given behavior, we might therefore expect less neural stability for the wet-dog shakes compared to the learned lever-pressing task. Alternatively, if sensorimotor circuits maintain a stable mapping to behavior more generally, we should expect single neuron activity patterns in MC and DLS to be stable in relation to the WDS behavior – albeit perhaps with less behaviorally modulated firing since these brain regions are dispensable for the behavior.

Given the stereotyped frequency of the WDS events, it is possible to identify them using an accelerometer attached to the head of each animal (Methods). Each WDS event lasted approximately 500 ms, and each animal performed on the order of 50 WDS per day. This allowed us to analyze them in a ‘trial-like’ manner, similar to our analyses of the lever-pressing task. We found that the accelerometer readouts corresponding to WDS events were consistent across trials over long time periods (Figure 6A), and we identified units in both DLS and MC whose activity was locked to the behavior. Consistent with the stable single neuron activity observed during the learned lever-pressing task, the neurons exhibited qualitatively similar firing patterns over time (Figure 6B), though there was weaker task modulation overall (Extended Data Figure 10). When computing PETH correlations over time, we also found that they remained stable throughout the period of recording (Figure 6C, Extended Data Figure 3B, 4B), although the baseline trial-to-trial similarity was lower than for the learned motor patterns associated with the timed lever-pressing task. These results are consistent with a lesser (or no) involvement of DLS and MC in the specification and control of WDS^75^. Hence, the observed activity patterns in MC and DLS during WDS, and consequently the high degree of stability over time, is likely to reflect the stability in the sensorimotor system as a whole, including in the behaviorally-locked activity of connected areas, which presumably process sensory feedback and motor efference^76^. This is also consistent with our finding of stable motor cortical activity in the lever-pressing task, where motor cortex is only necessary for learning but not for executing the task^6^.

**Figure 6:**
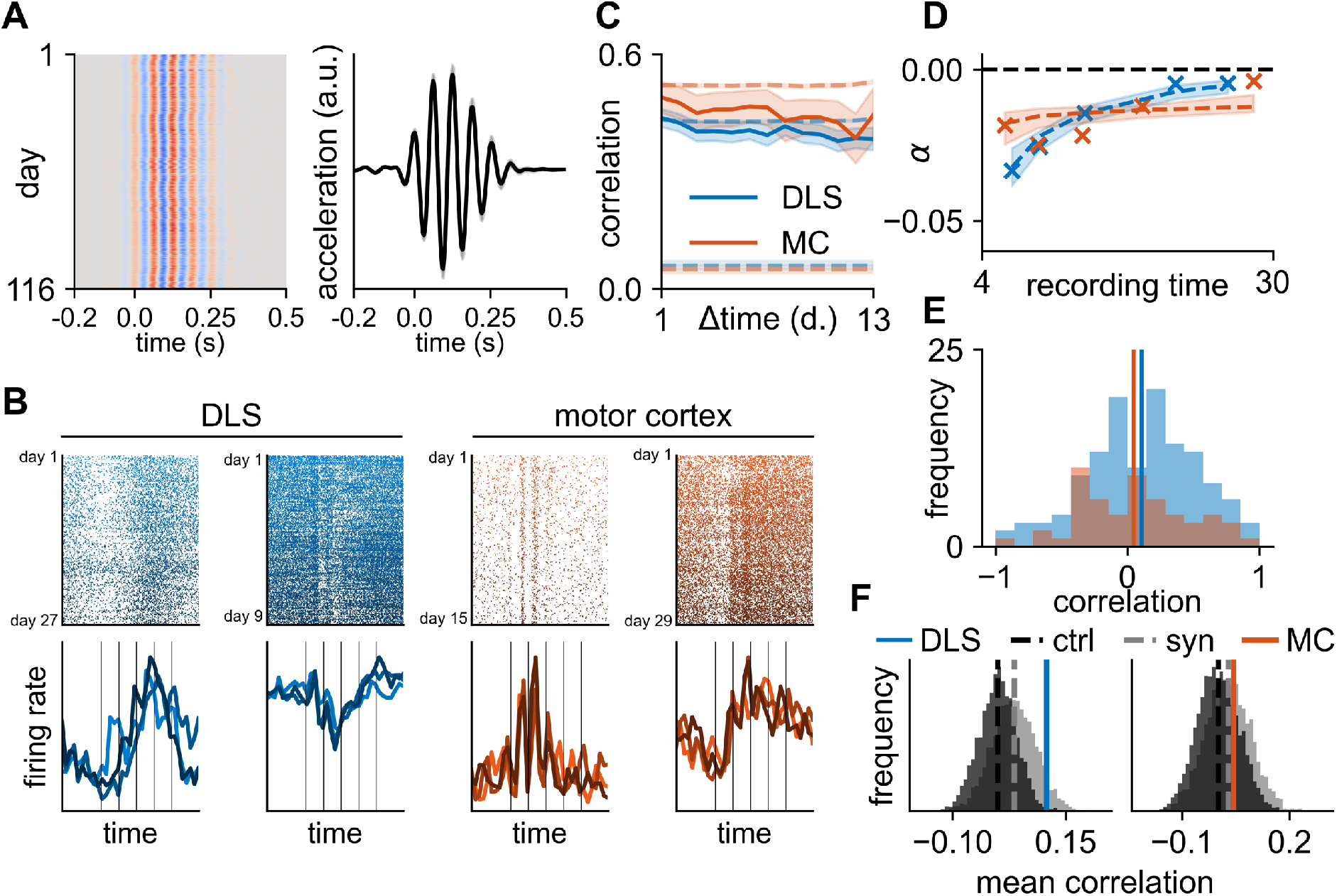
Neural activity is stable during an innate behavior (WDS). **(A; left)** Vertical acceleration across all 12,775 wet-dog shakes (WDSs) recorded over 116 days in an example animal. Each row corresponds to a single ‘trial’. **(A; right)** We computed the mean acceleration across trials for each day. Line and shading indicate the mean and standard deviation across all days as a function of time-within-trial. All kinematics and spike times were linearly time-warped to the median WDS frequency for each animal (Methods; warping coefficient = 1.01 ± 0.07; see Extended Data Figure 7 for results without time-warping). **(B; top)** Raster plots for two example units in DLS (left) and MC (right), illustrating units with firing patterns that are time-locked to the behavior over timescales of days to weeks. Color indicates the progression of time from day 1 (light) to the last day of recording (dark). **(B; bottom)** PETHs computed on three different days (early/middle/late) for each of the four example units. Vertical lines indicate peaks in the accelerometer trace. **(C)** Mean value of the correlation between PETHs calculated on separate days, averaged over all units recorded for at least 14 days in MC (red; n = 17) or DLS (blue; n = 66) and plotted as a function of time difference (see Extended Data Figure 4B for other recording thresholds). Shadings indicate standard error across units. Dashed lines indicate the similarity between non-identical neurons (lower) and in a resampled dataset with neural activity drawn from a stationary distribution (upper; Methods). **(D)** Rolling median of the stability index (crosses) for units with different recording durations. Bins are overlapping, with each neuron occurring in two bins (see Extended Data Figure 6D for the non-binned data). Dashed lines indicate exponential model fits to the non-binned data, and shadings indicate interquartile intervals from bootstrapping the units included in the model fits (Methods). **(E)** Distribution of the correlation between neural similarity and behavioral similarity on consecutive days for neurons recorded in DLS (blue) or MC (red). Vertical lines indicate average correlations. **(F)** Mean across units of the correlation between neural and behavioral similarity on consecutive days in DLS (blue; left panel) and MC (red; right panel). Dark grey histograms indicate control distributions from permuting the days in the behavioral data. Light grey histograms indicate the distributions of correlations in synthetic datasets where neural activity is determined entirely by behavior via a GLM.

To quantify the degree of stability for the population of recorded neurons during WDS, we computed stability indices for each neuron. Similar to our observations in the lever-pressing task, the stability indices were centered near zero, indicating largely stable circuits, but with a slow decay over time (DLS: α*_median_* = −0.014, τ*_median_* = 70 days, p < 0.001, n = 180 neurons; MC: α*_median_* = −0.016, *τ_median_* = 63 days, p < 0.001, n = 99 neurons; permutation tests). We expected that this apparent drift would be partly due to our finite recording durations as for the lever-pressing task. Consistent with this hypothesis, stability indices increased with recording duration in both DLS and MC (Figure 6D). Additionally, fitting an exponential model with a baseline to the average similarity across neurons suggested a decay to an asymptotic correlation of *γ* = 0.39 for DLS and *γ* = 0.29 for MC. These results show that the neural activity patterns associated with this innate behavior are stable over long timescales, similar to our observations for learned motor skills.

Based on our analyses of the lever-pressing task, we wondered whether some of the residual neural drift could be accounted for by changes in the kinematics associated with the WDS. We therefore investigated whether the motor output during WDS exhibited a systematic drift over time and found this to be the case for all animals (Extended Data Figure 8). To query whether the behavioral drift could be linked to the drift in neural activity, we computed the mean correlation between neural and behavioral drift on consecutive days. This analysis confirmed the presence of a weak but significant effect of behavioral drift on neural drift in DLS (Figure 6E; 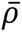 = 0.11, p = 0.005; permutation test). We also found a weak correlation in MC, although this did not reach the threshold for ‘statistical significance’ (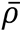 = 0.04, p = 0.24; permutation test). While these correlations are small, we found them to be consistent with the expectation from a synthetic model, where neural drift is entirely driven by behavioral drift (Figure 6F). This suggests that the small effect size could be due to the weak effect of the recorded neurons on the WDS behavior or the slow rate of drift in both behavior and neural activity, which contribute to a small signal-to-noise ratio. Both here and in the case of the timed lever-pressing task, we thus interpret the observed drift in neural activity to be accounted for, in large part, by slowly changing behavior.

## Discussion

We have investigated whether stereotyped motor behaviors are driven by stable single neuron dynamics (Figure 1) in two major nodes of the motor system that are involved in the acquisition of motor skills – MC and DLS. Using an RNN model, we first demonstrated the necessity of long-term single-unit recordings for answering this question (Figure 2). We then performed such recordings in rats trained to generate highly stereotyped task-specific movement patterns (Figure 3)^6, 59^. We found that the task-aligned activity of neurons in both MC and DLS was remarkably consistent over time, as expected for a stable control network. Recording single units for long durations was important to reveal this stability and distinguish it from constrained fluctuations on shorter timescales (Figure 4). We did observe a slow drift at the population level, which was accompanied by a concomitant drift in behavioral output (Figure 5). This is similar to previous reports of motor drift in expert performers^66^. Importantly, the drift in behavior was correlated with the recorded drift in neural activity, suggesting that the neural drift could be explained, in large part, by small but systematic behavioral changes. Finally, we showed that these observations extend to an innate behavior with trial-like structure (Figure 6), suggesting that stable sensorimotor circuits underlie stereotyped behavior, both learned and innate.

### Impact of behavioral variability in studies of neural stability

We showed how behavioral changes not fully constrained by the task can lead to the appearance of drift in single-unit neural activity patterns. This is particularly relevant since movements not relevant to the task are known to be strongly represented in cortical activity^49^. As a result, our reported neural stability in relation to both the learned and innate motor behaviors, and similar reports from other studies, should be seen as lower bounds on the neural stability associated with a hypothetical perfectly stable behavior. Additionally, this observation of correlated neural and behavioral drift highlights the importance of high-resolution behavioral measurements when investigating the stability of neural circuit dynamics, since most tasks studied in neuroscience do not fully constrain behavioral output^77^.

While the observed slow drift in neural and behavioral space in expert animals suggests that the changes in neural circuits occur in directions of neural state space that affect motor output, it remains to be seen whether this behavioral drift constitutes a learning process that optimizes a utility function such as energy expenditure^78^ or magnitude of the control signal^79^. Alternatively, it could reflect a random walk in a degenerate motor space that preserves task performance^25, 66^. Previous work has also suggested that motor variability could be explicitly modulated to balance exploration and exploitation as a function of past performance and task uncertainty^80^. If the behavioral drift we observe experimentally reflects such deliberate motor exploration, we might expect neural drift to be biased towards behaviorally potent dimensions to drive the necessary behavioral variability^81^. Conversely, if the behavioral drift is a consequence of inevitable drift at the level of neural circuits, neural drift might be unbiased or even preferentially target behavioral null dimensions to minimize the impact on task performance. Future studies will be needed to arbitrate between these possibilities.

### Prior studies of neural stability

It is worth noting the contrast between our results and previous studies that found task-associated neural activity in sensory and motor circuits to drift over time^18, 24, 35, 38, 56^. Some of these differences could reflect physiological differences between species, circuits, or cell function^31^, with recent studies e.g. showing differential stability between hippocampal cells representing time and odor identity in an odor discrimination task^82^. However, they could also reflect differences in methodology. For example, brain computer interfaces^35^ circumvent the natural readout mechanism of the brain, which could affect the stability of learned representations. Additionally, different statistical assessments of stability have previously been suggested to underlie discrepancies in the apparent neural stability underlying a primate reaching task^33^. Similarly, we find that accounting for the bias arising from finite recording durations is necessary to reveal the stability of sensorimotor circuits, and that unaccounted behavioral variability can confound analyses of representational drift in neural circuits. Furthermore, electrophysiology and calcium imaging can provide contrasting views on stability as discussed elsewhere^59, 83^. For the behaviors we probed in this study, electrophysiological recordings were essential to resolve neural dynamics on timescales of tens to hundreds of milliseconds^84^. Moreover, being able to record continuously over many weeks mitigates the need to stitch together separate recording sessions with potential movement of the recording electrodes and changes of spike waveforms between sessions^56, 59, 83^. We therefore expect that our understanding of neural stability will benefit further from recent impressive advances in recording technology^85–87^, especially if such advances can eventually be combined with methods for chronic recordings to track changes in the waveforms of individual neurons^59^.

The finding of stable neural correlates of motor output by us and others^29, 32, 34, 37^ can also be contrasted with recent work suggesting that neural activity patterns in posterior parietal cortex (PPC) change over a few days to weeks during a virtual navigation task with stable performance^15^. This discrepancy could arise from differences in methodology, recording duration, or limited behavioral constraints as discussed above. It could also reflect the fact that higher cortical regions are more sensitive to internal or external latent processes that lead to the appearance of drift due to an unconstrained environment. However, an alternative explanation is that higher-order brain regions, such as PPC or prefrontal cortex, accommodate drifting representations to allow fast learning processes or context-dependent gating of stable downstream dynamics^26, 88–90^. This is consistent with theoretical work on stable readouts from drifting neural codes^26, 31^, with our results supporting the hypothesis that stable representations are more likely closer to the motor periphery. These ideas are also consistent with a recent hypothesis in the olfactory domain that piriform cortex implements a ‘fast’ learning process with drifting representations, which drives a ‘slow’ learning process of stable downstream representations^18^. In the context of brain-computer interfaces, where a stable mapping between measured activity and system output is desirable^91^, these considerations suggest that decoding activity from motor cortex or even subcortical brain regions is preferable to higher-order cortical areas such as prefrontal cortex or PPC. Finally, our results provide experimental evidence that the stable motor cortical latent dynamics observed in previous work^43^ may be a consequence of dynamics that are stable at the level of single neurons.

### Maintaining stability in the face of dynamic network changes

Our findings of long-term stability in both MC and DLS raise questions of how this is achieved mechanistically and whether there are active processes maintaining stability of network dynamics. Manipulation studies in both motor and sensory circuits suggest that such processes do exist in the case of large-scale perturbations. For example, it has previously been shown that motor circuits can recover their activity and function after invasive circuit manipulations by returning to a homeostatic set-point, even in the absence of further practice^92^. At the single neuron level, there are also intrinsic mechanisms keeping the firing rates of neurons in a tight range. Indeed, an increase in the excitability of individual neurons has been observed following sensory deprivation in both barrel cortex^93^ and V1^94–96^, with V1 also recovering higher-order network statistics^97^. These observations suggest that the brain uses homeostatic mechanisms to overcome such direct perturbations. Of course, these perturbations are large and non-specific compared to the changes that occur during normal motor learning, which instead consist of gradual synaptic turnover and plasticity. However, it is plausible that some of the same mechanisms that help restabilize networks following large-scale perturbations could also be involved in maintaining network stability under natural conditions^98, 99^.

Taken together, our results resolve a long-standing question in neuroscience by showing that the single neuron dynamics associated with stereotyped behaviors, both learned and innate, are stable over long timescales. However, they raise another mechanistic question of how new behaviors are learned without interfering with existing dynamics – that is, how does the brain combine long-term single-unit stability with life-long flexibility and adaptability^13, 27, 28,100–102^? This is an essential yet unanswered question for neuroscience, and future work in this area will likely require more elaborate experimental protocols combining interleaved learning of multiple tasks with long-term neural recordings and high-resolution behavioral tracking to elucidate the mechanistic underpinnings of network stability and flexibility.

## Acknowledgements

We are grateful to Kiah Hardcastle, Cengiz Pehlevan, Ta-Chu Kao, Guillaume Hennequin, and Marine Schimel for their feedback on the manuscript. This work was supported by a Gates Cambridge scholarship and Nordea-fonden (KTJ); a Helen Hay Whitney postdoctoral fellowship, the Zuckerman STEM Leadership Program postdoctoral fellowship, and the Women in Science Weizmann Institute of Science Award (NKH); a Life Sciences Research Foundation and Charles A. Kings Foundation postdoctoral fellowship (AKD); an EMBO postdoctoral fellowship ALTF1561-2013 and an HFSP postdoctoral fellowship LT 000514/2014 (SBEW); and NIH grants R01-NS099323-01 and R01-NS105349 (BPÖ).

## Methods

### Data Analysis

#### Animal training and data acquisition

Female Long Evans rats (n = 6) were trained in an automated home-cage system on a lever-pressing task as described previously^6, 58, 59^. In short, animals were rewarded for pressing a lever twice with an inter-press interval of 700 ms. Electrophysiological data was recorded from layer 5 of motor cortex (MC; n = 3) and from dorsolateral striatum (DLS; n = 3) and spike-sorted as described by Dhawale et al.^59^. Data from all animals have previously been used by Dhawale et al.^57^. The care and experimental manipulation of all animals were reviewed and approved by the Harvard Institutional Animal Care and Use Committee.

#### Behavioral tracking

Videos were recorded at 120 Hz during the lever-pressing task from two cameras positioned at the left and right side of the home cage relative to the lever. Automated behavioral tracking was carried out using DeeperCut^62, 63^. For each of 500 frames from each camera, the corresponding forelimb of the animal was manually labelled. This was used as a training dataset for DeeperCut to generate full trajectories for all trials followed by interpolation with a cubic spline. Clustering of task trials based on behavioral readouts was carried out using forelimb positions tracked by DeeperCut as well as accelerometer data from an accelerometer attached to the skull of each animal. These features were embedded in a t-SNE space and clustered using density-based clustering^103^. Only trials falling in the largest cluster for each animal (range of 37% to 93% of trials across animals) and with inter-press intervals (IPIs) between 600 ms and 800 ms were included in the analyses to minimize behavioral variability.

#### Detection and classification of wet dog shakes

To identify wet dog shakes (WDS), accelerometer data was first passed through a 12-20 Hz filter and the magnitude of the response calculated as 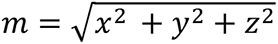. A moving average of *m* was calculated with a window size of 1/6 seconds, and WDS events identified as periods with *m* > 0.03. Peaks were found in a window of 800 ms centered at the middle of each WDS event and identified as local maxima or minima with a prominence of at least 0.07 times the difference between the highest maximum and lowest minimum in each channel. WDS events were aligned to the first positive peak in the vertical (z) channel and time-warped according to the inter-peak separation in this channel. Aligning to either horizontal channel gave similar results, and the vertical channel was preferred to avoid the degeneracy of the horizontal plane.

#### Time-warping

For all analyses of experimental data, we time-warped neural activity and behavior using piecewise linear warping^104^ with parameters that aligned the two lever presses across all trials (see Extended Data Figure 7 for analyses without time-warping). We did this since neurons in DLS and MC have previously been shown to have activity patterns linked to these events^59^. Time-warping of spike data in the lever-pressing task was carried out by linearly scaling all spike times between the first and second presses by a factor 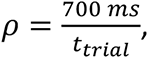 where *t_trial_* is the inter-press interval (IPI) in a given trial. All spike times after the second press were shifted by 700 *ms* − *t_trial_*. Warping of behavioral data was carried out by fitting a cubic spline to the trajectories and extracting time points at a frequency of 120 Hz prior to the first press, *ρ* × 120 Hz between the two presses, and 120 Hz after the second press. The warping coefficient *ρ* had a mean of 1.00 and a standard deviation of 0.07 across all trials and animals.

Warping of spike data for the wet dog shakes was carried out by linearly scaling all spike times between a quarter period before the first peak (*t*_1_) and a quarter period after the last peak (*t*_2_) by a factor 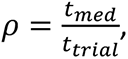 where *t_trial_* is the period of the oscillation in a given trial and *t*_&’(_is the median period across all trials and sessions for a given animal. All spike times before *t*_1_ were shifted by *t*_1_ × (*ρ* − 1) and all spike times after *t*_/_ were shifted by *t*_2_ × (*ρ* − 1). Warping of behavioral data was carried out by fitting a cubic spline to the accelerometer data and extracting time points at a frequency of 300 Hz prior to *t*_1_, *ρ* × 300 Hz between *t*_1_ and *t*_2_, and 300 Hz after *t*_2_. The first detected positive peak was assigned a time of zero for each WDS. The warping coefficient *ρ* had a mean of 1.01 and a standard deviation of 0.07 across all trials and animals.

Data between 0.1 seconds before the first tap and 0.1 after the second tap was used for all analyses of the lever-pressing task, and data between 0.2 seconds before and 0.5 seconds after the first accelerometer peak was used for all WDS analyses.

#### Similarity of neural activity

PETHs were calculated for each session by summing the spikes across all trials for each time-within-trial. We convolved the resulting spike counts with a 15 ms Gaussian filter for the lever-pressing task, and with a 10 ms Gaussian filter for the WDS behavior. Pairwise PETH similarities between sessions were calculated as the Pearson correlation between **u** and **v**, where **u** and **v** are vectors containing the PETHs at 20 ms resolution. PETHs were normalized by z-scoring for visualization in Figure 4A for each unit, and by total spike count on each day for the PETHs in Figure 4B and 6B. Neural similarity as a function of time difference was calculated by computing the pairwise similarity of the PETHs for each unit across every pair of days in which the PETH contained at least 10 spikes. The pairwise similarities for each time difference were averaged across units in Figures 4C and 6C, after first averaging over all PETH pairs separated by the same time difference for each individual unit.

We restricted all analyses to neurons that were ‘task-modulated’. To define task-modulation, we computed a PETH for odd and even trials separately for each recording day and considered the correlation between this pair of PETHs on each day. We then averaged the result across days for each neuron. A neuron was considered task-modulated if this measure of same-day similarity exceeded *ρ*_0_ = 0.15. This resulted in 221 of 363 neuron being task-modulated in DLS during lever-pressing task, 446 of 795 in MC during the lever-pressing task, 344 of 1250 in DLS during the WDS behavior, and 372 of 904 in MC during the WDS behavior.

#### Control analyses for stability as a function of time difference

In Figures 4C and 6C, we include a positive and a negative control for the neural similarity as a function of time difference. Here we provide a description of how these were computed. For the negative control, we computed the similarity between non-identical pairs of neurons. This can be seen as the asymptotic similarity in the limit of complete neural turnover but with constant population statistics (i.e. each neuron corresponds to a randomly sampled neuron from the population). This similarity was averaged across 1000 pairs of randomly sampled neurons, with each pair being recorded in a single animal. For the positive control, we resampled the activity of each neuron on all trials in which it was recorded, with replacement, from the total distribution of recorded trials across days. We then computed the similarity as a function of time difference for all neurons as in the original data. The process of resampling and computing similarity was repeated 100 times, and the figures indicate mean and standard error across samples. This control thus corresponds to the hypothetical similarity in the case where all neurons have a fixed distribution over firing patterns (i.e. neural activity is stable), and where the global distribution of firing patterns is matched to the data.

#### Alignment of neural dynamics

Aligning neural dynamics using CCA requires simultaneous recording of a large number of neurons. Since our recordings were asynchronous, this was in general not the case (c.f. Figure 3C). For this analysis, we therefore focused on a smaller subset of the data, where 16 neurons were simultaneously recorded for a week and fired at least 10 spikes on each day. This dataset corresponds to days 8-14 of the DLS animal indicated in Figure 3C. We first computed the ‘single neuron similarity’ in this dataset by computing the average PETH correlation across all neurons for each pair of days. We then computed the mean and standard deviation of this measure across all pairs of days separated by the same time difference. This provided a measure of the similarity of neural dynamics in a constant coordinate system with the axes aligned to individual neurons. For comparison with this, we also computed the similarity of neural activity when aligning the neural dynamics using CCA for each pair of sessions. To do this, we followed the approach outlined by Gallego et al.^43^ to align the dynamics on day ‘b’ to the dynamics on day ‘a’ across all pairs of days. This alignment was carried out at the level of PETHs rather than single trials. For these analyses, we aligned the dynamics across all neurons and considered the average correlation across all the resulting dimensions (i.e. the similarity was the average of all CCs). This addresses the question of whether stability increases if we allow for linear transformations of the coordinate system in which we characterize neural dynamics.

#### Exponential model fits and stability indices

To assess the stability of neural activity over time, we examined the Pearson correlation *ρ* between the computed PETHs as a function of the time difference *δt* between PETHs. We then fitted an exponential model 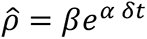 to this data for each neuron recorded for at least 4 days. This was done to better quantify the putative drift in neural activity across neurons by learning a parameter *α* that encompasses the rate of drift for each neuron. Here, *β* is an intercept describing the expected similarity for two sets of trials recorded on the same day, and *α* determines the rate of change of neural similarity. For this fit, we constrained *β* to be between −1 and +1 by passing it through a tanh transfer function, since Pearson correlations must fall in this interval. The parameters were optimized to minimize the squared error between the predicted (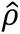) and observed (*ρ*) PETH correlations. This was done numerically, and the optimization was initialized from a linear fit to the data 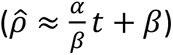. We denote the learned parameter *α* with units of inverse time as a ‘stability index’. This is related to the time constant of an exponential decay model via *α* = −*τ*^-1^, with the fitting of *α* being numerically more stable as it avoids *τ* approaching infinite values for slow decays. All data points with a time difference of at least 1 day were used to fit the models. The mean error of the model fit was quantified for each neuron as 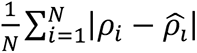, where |⋅| indicates the absolute value, and the sum runs over all *N* data points (Extended Data Figure 6B). Significance of median stability indices being different from zero was calculated by shuffling the vector of time differences for each unit 2,000 times, each time computing the median of the stability indices across all units and counting the fraction of shuffles where the median stability index was smaller than the experimentally observed median.

For comparison with this single-timescale model, we also considered a model which decayed to a learnable baseline *γ*: 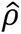 = *βe^αδt^* + *γ*. We did this since the presence of constrained latent processes could lead to a decay in neural similarity to a non-zero asymptotic value at long time differences. Clearly, the single-timescale exponential decay arises as a special case of this model for *γ* = 0. However, it is also worth noting that a linear model, commonly used in the literature^18, 24, 38, 43, 56, 59^, arises as *γ* → −∞. Intuitively, this is the case since any finite region of 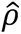 is in the initial linear regime of an exponential that decays to −∞. This model with a baseline thus serves as a generalization of both the linear and exponential models. When fitted to the neuron-averaged data and evaluated using hold-one-out crossvalidation, this model performed comparably to or better than the simple exponential decay model on all four datasets (recordings from DLS/MC across the two tasks). Additionally, using this same crossvalidated evaluation metric, the exponential model consistently outperformed a linear model, suggesting that this is a more appropriate single-timescale model.

#### Stability as a function of recording duration

To extrapolate our stability indices to long recording durations across the population, we fitted a model to the stability index *α* as a function of recording time *T* of the form 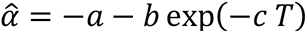. We fitted the model by minimizing the L1 error between the observations and model fit across neurons 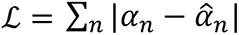 and restricted all parameters {*a*, *b*, *c*} to be positive. In this model, the asymptotic stability is given by 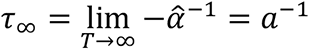. To construct confidence intervals for this analysis, we subsampled the data points for each neuron {*α_n_*, *T_n_*} with replacement and repeated the model fitting procedure. Results are reported as medians and interquartile ranges by considering the 25^th^, 50^th^ and 75^th^ percentile of the corresponding distribution over *τ*_∞_. While the model itself was fitted to the raw data, we denoised the data for the visualization in Figure 4E by plotting the median stability index across neurons binned by recording duration. The bins were selected with partial overlap (each neuron occurred in two bins), and the x-value indicated for each data point in the figure is the average recording duration for each neuron in the corresponding bin.

To compute the stability as a function of subsampled recording duration in Figure 4F, we used successive maximum time differences from **δ*t_max_* = 3 to **δ*t_max_* = 13 days. We then considered the average similarity as a function of time difference in Figure 4C, using only data up to and including **δ*t_max_*. We computed stability indices for these subsets of data as described above and plotted the stability as a function of **δ*t_max_*.

#### Behavioral similarity

To compute behavioral similarity as a function of time difference, we first extracted instantaneous velocities of both forelimbs in the vertical and horizontal directions as the first derivative of the time-warped cubic spline fitted to position as a function of time. We computed the pairwise behavioral similarity between sessions as the correlation between the mean velocity profiles across all trials from the corresponding sessions, averaged across both forelimbs and the vertical/horizontal directions.

To compute the correlation between neural and behavioral drift rates, we considered the behavioral similarity on pairs of consecutive days together with the neural similarity across the corresponding days, quantified using PETH correlations as described above. We then considered the distribution of neural and behavioral similarities across all pairs of consecutive days for each recorded unit and computed the correlation between these two quantities. Finally, we computed the mean of this correlation across the population of units recorded from either DLS or MC. As a control, we permuted the behavioral data across days to break any correlations between the neural and behavioral drift rates and repeated the analysis. In Figure 5E and 6F, null distributions are provided across 5,000 such random permutations. For these analyses, we did not include the first day of recording for any unit since this data was used to fit the synthetic control data (see below). Furthermore, we only considered neurons with at least 4 pairs of consecutive recording days (after discarding the first day of recording), such that all correlations were computed on the basis of at least 4 data points.

#### Stability of population decoding

To investigate the stability of a population decoder, we considered the same week-long subset of data as for the alignment of neural dynamics. We first square root transformed the neural data and convolved it with a 40 ms Gaussian filter, similar to previous work. We then trained a crossvalidated ridge regression model to predict the left and right forelimb trajectories from neural activity using data from each single day and tested the model on all other days. Finally, we computed the performance of this decoder as a function of time difference between testing and training. For all decoding, we offset behavior from neural activity by 100 ms to account for the fact that neural activity precedes kinematics, similar to previous work in primates^23, 105^. To test whether the decoder exhibited a significant decrease in performance as a function of time difference, we performed a bootstrap analysis by resampling with replacement the similarity as a function of time difference (i.e. we resampled ‘pairs of days’) and computing the slope of a linear fit to the data.

For comparison with this decoding model, we also considered decoding performance in an aligned latent space. To do this, we again considered all pairs of days and matched the number of trials on each pair of days to facilitate alignment (i.e. we discarded the later trials on the day with most trials). We then used PCA to reduce the dimensionality of the data from 16 to 10 for each day and trained our crossvalidated ridge regression model to predict behavior from this latent neural activity on the training data. At test time, we aligned the PCs on the test day to the PCs on the training day and predicted behavior from these aligned PCs. This follows the procedure described in previous work^43^. Note that alignment was in this case done at the level of single trials rather than trial-averaged PETHs. Finally, we considered the decoding performance as a function of time difference for this aligned decoder.

#### GLM model fitting and analysis

To investigate the correlation between neural and behavioral drift rates in synthetic data, where neural drift is determined entirely by behavioral drift (Figure 5E), we first fitted a linear-nonlinear Poisson GLM to the first day of recording for each neuron. This model took the form ***y***_*t* = 1_ ∼ *Poisson*(exp[***Wx***_*t* = 1_]), where ***y***_#_ are the observed spike counts on day *t* across time bins (here a concatenation of trials and bins within each trial), ***x****_t_* is a set of input features, and ***W*** is a weight matrix that is learned by maximizing the log likelihood of the data. As input features, we used the velocity of both forelimbs in the x-y plane for the lever-pressing task, and the accelerometer readout in 3 dimensions for the WDS task. In both cases, we included a 200 ms window of kinematics surrounding each 20 ms bin of neural activity in the feature vector.

After fitting the model to data from day 1, we proceeded to generate synthetic neural activity by drawing spikes from the model 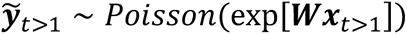 for all subsequent days using the recorded behavior ***x***. We then constructed PETHs for each unit and session, as described for the experimental data, and repeated the analysis correlating behavioral similarity with neural similarity on consecutive days for this synthetic dataset. We repeated the sampling and analysis process 5,000 times to generate a distribution of neural-behavioral correlations from this synthetic model and computed p values as the fraction of synthetic correlation values that were smaller than the experimentally observed value. When performing these analyses, we discarded the first day of recording in both the synthetic and experimental data since this was used to fit the GLM.

To test the stability of the encoding model (Extended Data Figure 5E), we computed predicted firing rates on all days not used for training. We then correlated the square root of the predicted firing rate with the square root of the observed spike count (for variance stabilization). Finally, we averaged this test correlation across all neurons that had a training correlation of at least 0.1 and were recorded for at least 7 days.

### Recurrent network modelling

#### Network architecture and training

The RNNs used in Figure 2 consisted of 250 recurrently connected units and 5 readouts units, which were simulated for 250 evenly spaced timesteps to generate 5 target outputs drawn from a Gaussian process with a squared exponential kernel that had a timescale of 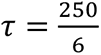. The RNN dynamics were given by

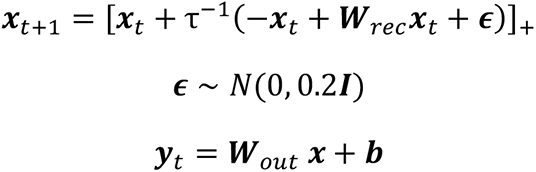

***W****_rec_*_’,_, ***W****_out_*, ***b***, and ***x***_0_were optimized using gradient descent with Adam (Kingma and Ba 2014) to minimize the loss function

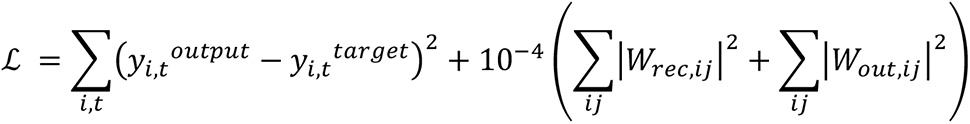

We used a learning rate of 0.0005 and batch size of 20 to train all networks.

#### Similarity measures

100 instances of each network were run to constitute a set of trials (a ‘session’). Observation noise was added to all neural activities *x* by drawing spikes from a Poisson noise model *s* ∼ *Poisson*(*λx*), where *λ* is a constant scaling factor for each session used to scale the mean activity to 6.25 Hz. PETHs were constructed by averaging the activity of each unit across all trials for a given network. PETH similarity was computed as the Pearson correlation between PETHs as for the experimental data. Behavioral similarity was computed as the mean RNN output correlation across pairs of trials for each pair of sessions. Latent similarity was computed by first convolving the single-trial activity with a 30 ms Gaussian filter. The activities of non-overlapping groups of 50 neurons were then concatenated into 50xT matrices for each session to simulate different simultaneously recorded populations of neurons. Here, T is the number of time bins per trial (250) times the number of trials per session (100). The 50xT matrices were reduced to 10xT matrices by PCA, and the resulting matrices were aligned by CCA across networks. The CCA similarity for a pair of networks and group of neurons was computed as the mean correlation of the top 4 CCs. This procedure was intended to mirror the analysis by Gallego et al.^43^.

#### Interpolating networks

To interpolate the networks in Figure 2D & 2E, two networks were first trained independently to produce the target output, generating two sets of parameters

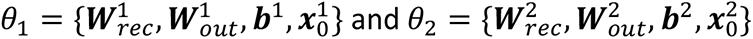

Seven new parameter sets *θ^dt^* were generated by linear interpolation between *θ*_1_ and (0.3*θ*_1_ + 0.7*θ*_2_), or equivalently by considering seven networks spanning the first part of a linear interpolation between *θ*_1_ and *θ*_2_. We chose not to consider the full interpolation series since neural activities became uncorrelated before the parameters were fully uncorrelated (Figure 2E), and we were interested in the range of parameters where neural activity drifted. For each interpolated parameter set, 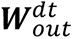 was fixed and the remaining parameters were finetuned on the original loss to ensure robust performance. Note that this procedure is merely used to generate a phenomenological model of a motor circuit with drifting connectivity and stable output, and it should not be interpreted as a mechanistic model. For the control network, the same interpolation and re-optimization procedure was carried out, but in this case interpolating between *θ*_1_and *θ*_1_(i.e. itself), such that the only differences between networks were fluctuations around the original connectivity. The whole procedure of training two initial networks and interpolating was repeated 10 times, and results in Figure 2E are reported as the mean and standard deviation across these repetitions.

## Extended Data figures

**Extended Data Figure 1:**
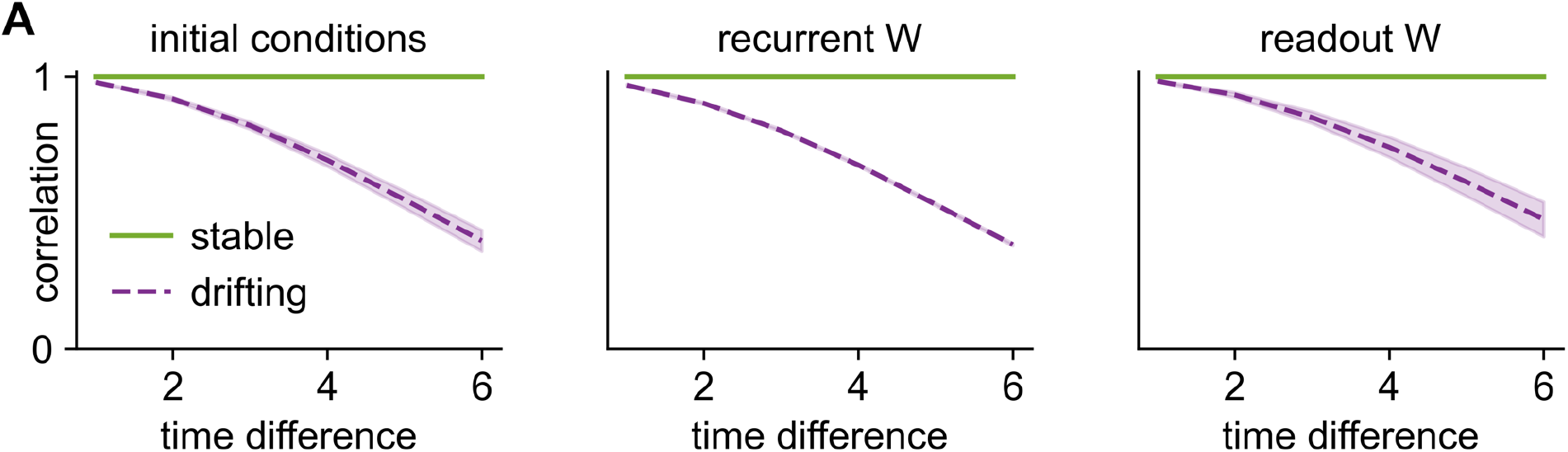
RNN parameter interpolation. **(A)** Mean correlation between the initial conditions (left), recurrent weight matrices (center), and readout weight matrices (right) of the simulated RNNs as a function of time difference for the stable and unstable networks. Shading indicates standard deviation across 10 repetitions of training and interpolation.

**Extended Data Figure 2:**
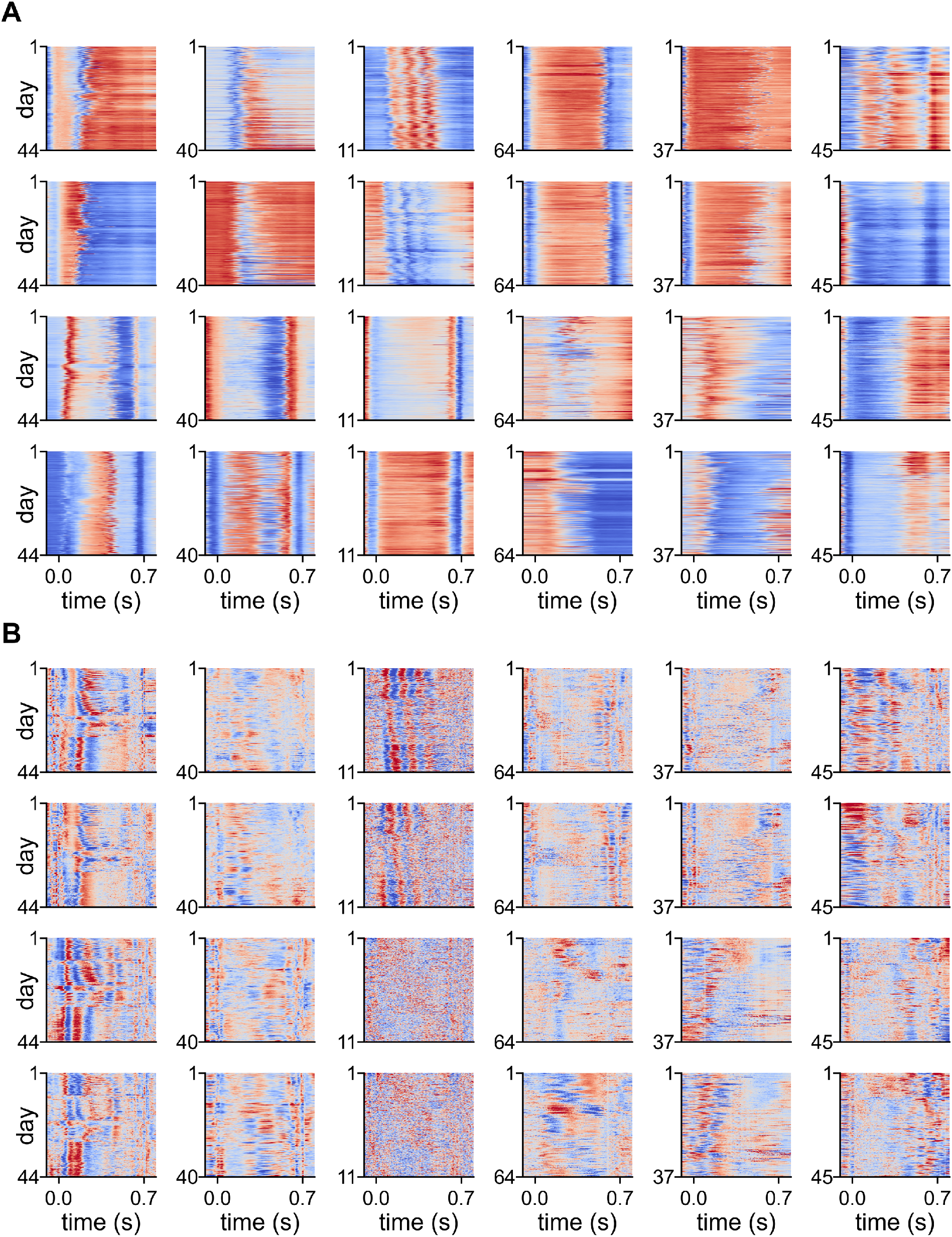
Kinematics of all animals. **(A)** Heatmaps showing the forelimb trajectories of each animal on every trial across all days. x-axes indicate time within trial and y-axes indicate trial number from first (top) to last (bottom). Each column corresponds to a single animal (first three: DLS, last three: MC). The rows illustrate the trajectories of the right forelimb parallel and perpendicular to the floor, followed by the left forelimb parallel and perpendicular to the floor. The second animal from the left corresponds to the example used in Figures 3A, 5A and 5B. **(B)** Heatmaps showing the z-scored velocity of each animal on every trial across all days for the animals in (A). The rows illustrate the velocity of the right forelimb parallel and perpendicular to the floor followed by the left forelimb parallel and perpendicular to the floor.

**Extended Data Figure 3:**
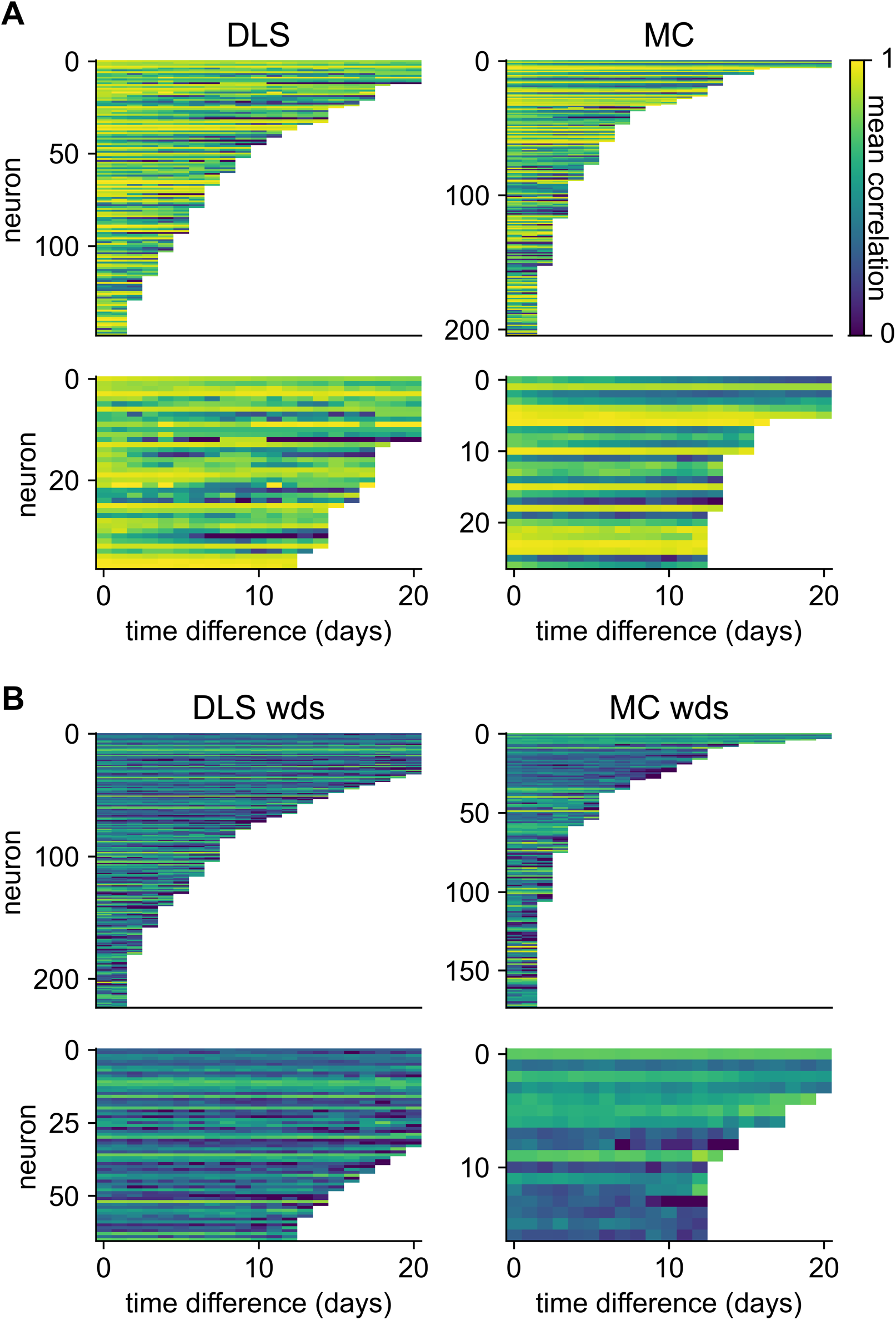
Similarity as a function of time difference for all neurons. We computed the PETH correlation as a function of time difference for all neurons, taking the average across all pairs of days separated by the same time difference for each neuron. This figure shows the average similarity as a function of time difference for neurons recorded in DLS (left) or MC (right) during the lever-pressing task **(A)** and the wet-dog shake behavior **(B)**. Upper panels indicate all neurons recorded for at least 3 days, lower panels indicate neurons which were recorded for at least 14 days and therefore included in Figure 4C & 6C. Neurons were sorted by recording duration.

**Extended Data Figure 4:**
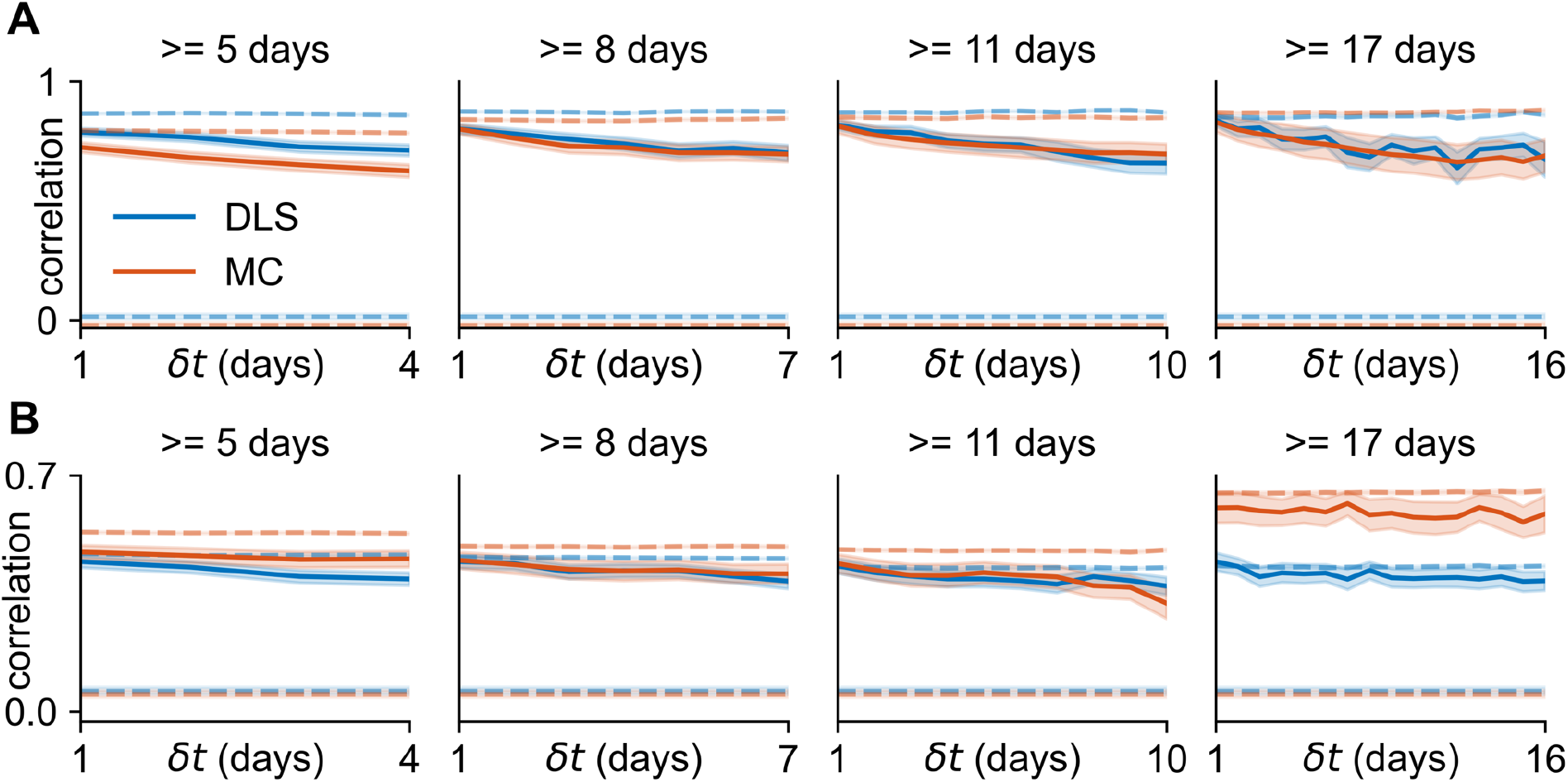
Stability as a function of time difference for different recording durations. **(A)** We performed analyses as in Figure 4C, plotting the neural similarity as a function of time difference for neurons recorded for at least N days, with N ranging from 5 to 17 (c.f. N = 14 in Figure 4C). Dashed lines indicate controls as in Figure 4C. **(B)** As in (A), now for the wet-dog shake behavior instead of the lever-pressing task (c.f. Figure 6C).

**Extended Data Figure 5:**
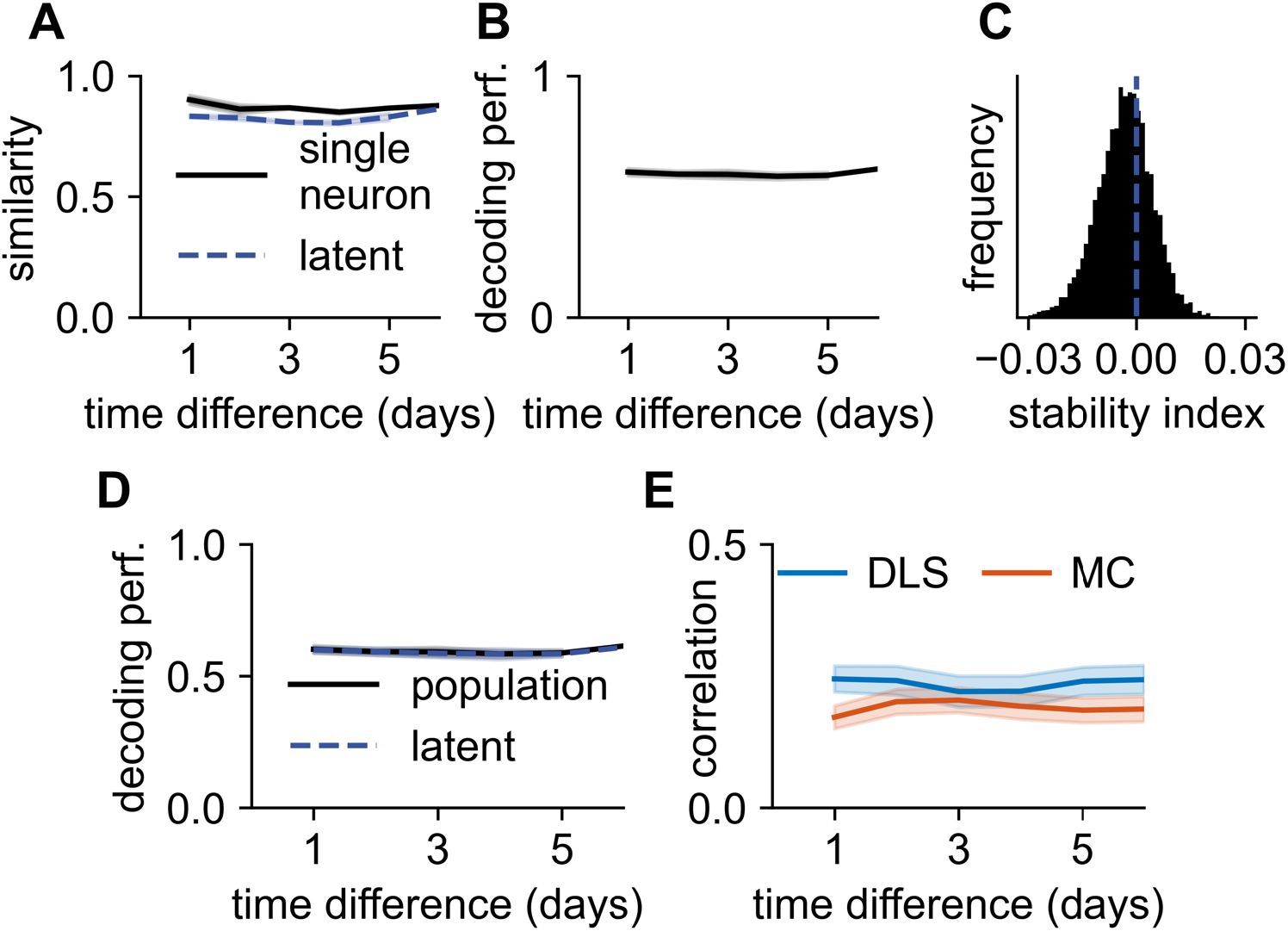
Latent stability and neural decoding. It has previously been reported that stable neural activity can be identified in a common latent space even when there is a turnover of recorded neurons^43^. As we show in Figure 2E, this can be consistent with either stable or drifting single unit activity. While we have already shown a high degree of similarity for single neurons, here we investigate whether ‘aligning’ the neural activity between sessions can identify a common subspace with even higher similarity. These analyses require simultaneous recording of a large population of neurons, which in general was not the case for our dataset (c.f. Figure 3C). Instead, we considered a single week of recording in a single animal with recordings from DLS (day 8-14 in Figure 3C), where we simultaneously recorded 16 neurons firing at least 10 spikes during the task on each day. **(A)** We first computed the similarity as a function of time difference as the correlation between single neuron PETHs, averaged across neurons (black line). We then proceeded to align the neural activity on each pair of days using CCA and computed the similarity in the resulting aligned space as the average correlation across all dimensions. This CCA-aligned similarity was generally lower than the similarity averaged over individual neurons, suggesting that the neuron-aligned coordinate system is more stable than the CCA-aligned alternative (note that CCA performs a greedy alignment rather than finding the optimal alignment, which would provide an upper bound on the single neuron similarity). **(B)** We proceeded to consider population decoding of behavior from neural activity, using the same data as in (A). We fitted a linear model to predict the trajectories of the left and right forelimbs from neural activity on each day using crossvalidated ridge regression, and we tested the models on data from all other days. Here, we plot the performance as a function of time difference, averaged across the vertical and horizontal directions and both forelimbs. Line and shading indicate mean and standard error across pairs of days with a given time difference. **(C)** We proceeded to compute stability indices for the data in (B) to see whether there was a significant negative trend. We bootstrapped the individual datapoints (before taking the mean) 10,000 times and estimated stability indices from each surrogate dataset. The distribution over the resulting stability indices was not significantly different from 0 (p = 0.34). **(D)** While the analysis in (A) suggests that the single neurons provide a good coordinate system for stable representations, it does not address the question of whether an aligned low-dimensional manifold can provide better decoding^43^. We therefore proceeded to train a population decoding model as in (B), but where the decoder was trained on the top 10 PCs from a single day and tested on the top 10 PCs from every other day after alignment via CCA^43^ (blue dashed line). We found that decoding performance from this aligned latent space was almost identical to the decoding performance from raw neural activity (black line). This provides further evidence that the stable aligned dynamics identified in previous work are the result of stable single-unit tuning curves. **(E)** Finally, we considered how the relationship between kinematics and neural activity changed over time at a single neuron level. We used the GLM discussed in Figure 5E to predict neural activity from behavior. This GLM was trained on the first day of recording for each neuron and tested on each subsequent day. The figure shows the correlation between the predicted firing rate and true spike count as a function of time difference, averaged across all neurons which were recorded for at least a week and had a training correlation of at least 0.1. Blue indicates neurons recorded from DLS, red from MC, and shadings indicate standard errors across neurons.

**Extended Data Figure 6:**
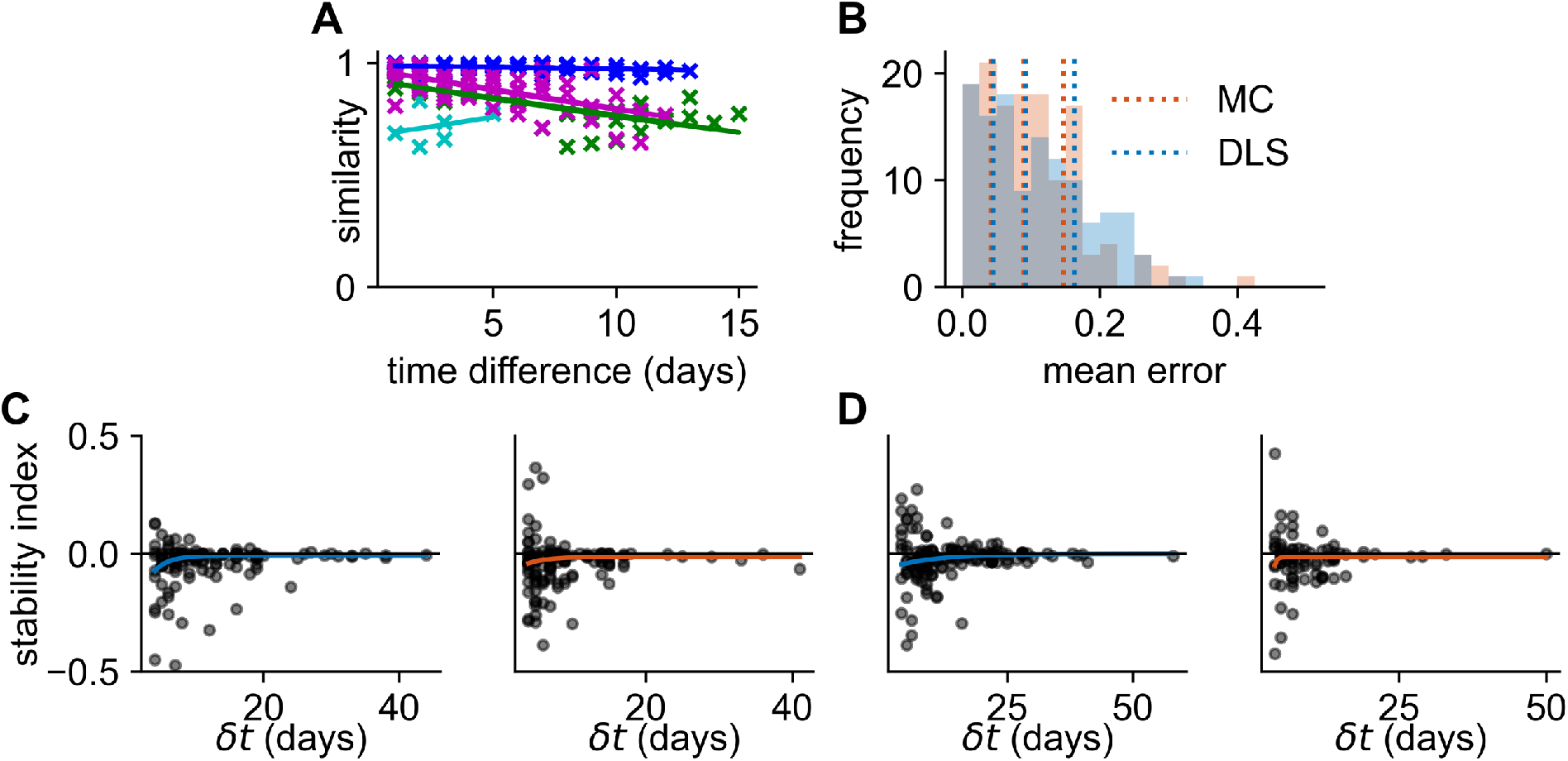
Exponential model fits and stability indices. **(A)** Plots of PETH similarity against time difference for four example units (colors) together with exponential fits illustrating a range of different decay rates, baseline similarities, and durations of recording. Note that one of these example units (cyan) exhibits an apparent increase in stability over time due to the noisy nature of the data. Indeed, in a perfectly stable model (such as the stable RNN in Figure 2E), neurons will be as likely to exhibit such an increase as they are to exhibit a decrease in similarity over time, leading to a median stability index of 0. Such noise is mitigated by increasing recording durations. **(B)** Distribution of the mean error of each model fit across the population of neurons recorded from MC (red) or DLS (blue). Vertical dashed lines indicate quartiles of the distributions. **(C)** Stability indices for all neurons recorded from DLS (left; blue) or MC (right; red) during the lever-pressing task. Solid lines indicate exponential fits as in Figure 4F. As the time difference increases, the variance decreases (due to the increase in data), and the median stability index gradually increases (c.f. solid lines). **(D)** As in (C), for the WDS behavior.

**Extended Data Figure 7:**
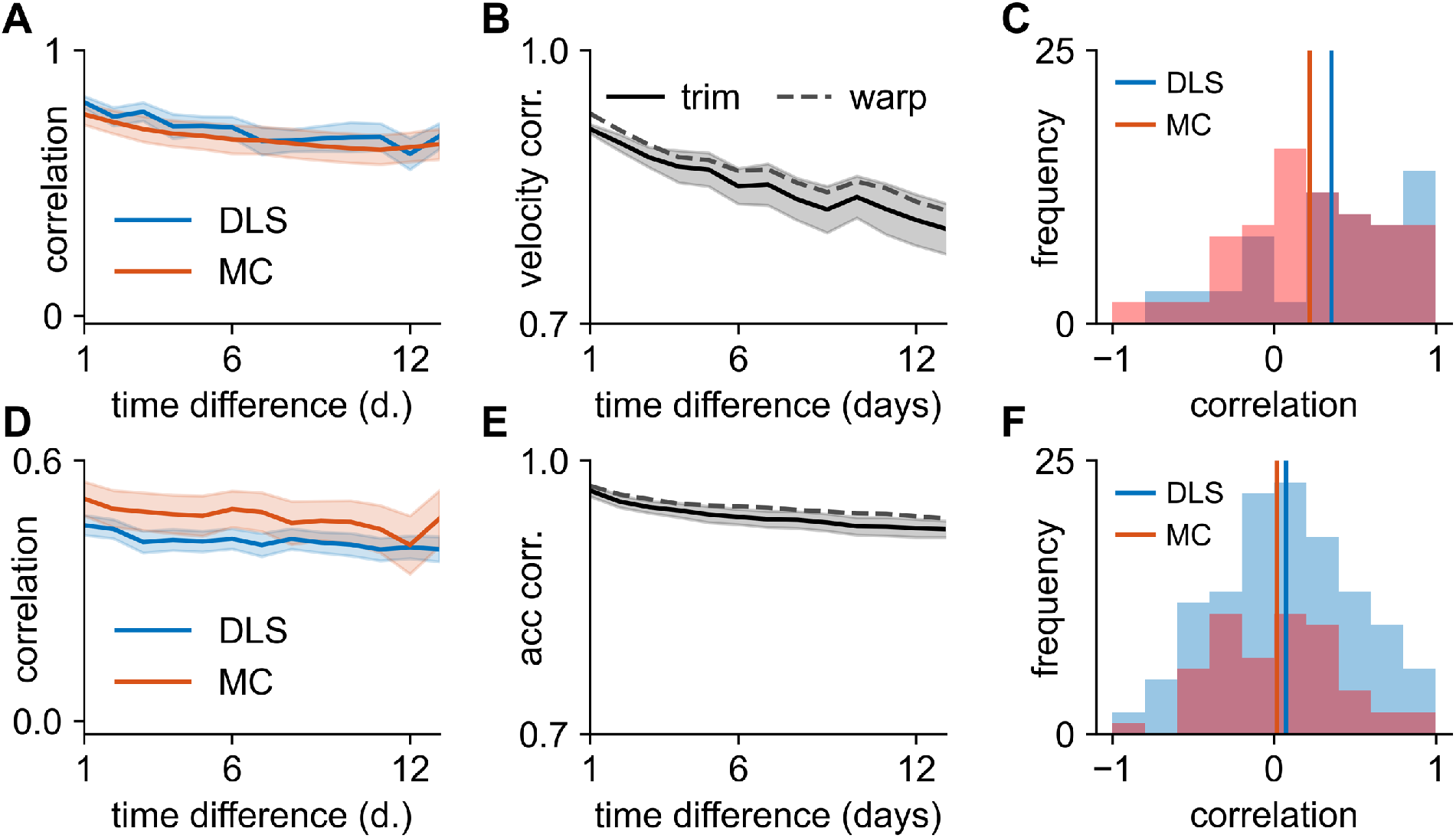
Results are not dependent on time-warping. In this figure, we reproduce some of the key analyses of the paper after aligning trials by ‘trimming’ to a fixed duration rather than the time-warping used in the main text. **(A)** Neural similarity as a function of time difference for neurons recorded for at least 14 days in the lever-pressing task in either DLS (blue) or motor cortex (red). Note similarity with Figure 4C using time-warping. **(B)** Kinematic similarity in the lever-pressing task as a function of time difference across all animals. Solid line and shading indicate mean and standard error across animals after trimming. Dashed line indicates the mean after time-warping. Note that time-warping better aligns kinematics, which is the primary motivation for its use in the main text. **(C)** correlation between neural similarity and kinematic similarity on consecutive days (c.f. Figure 5D). **(D-F)** As in (A-C), now for the wet-dog shake behavior.

**Extended Data Figure 8:**
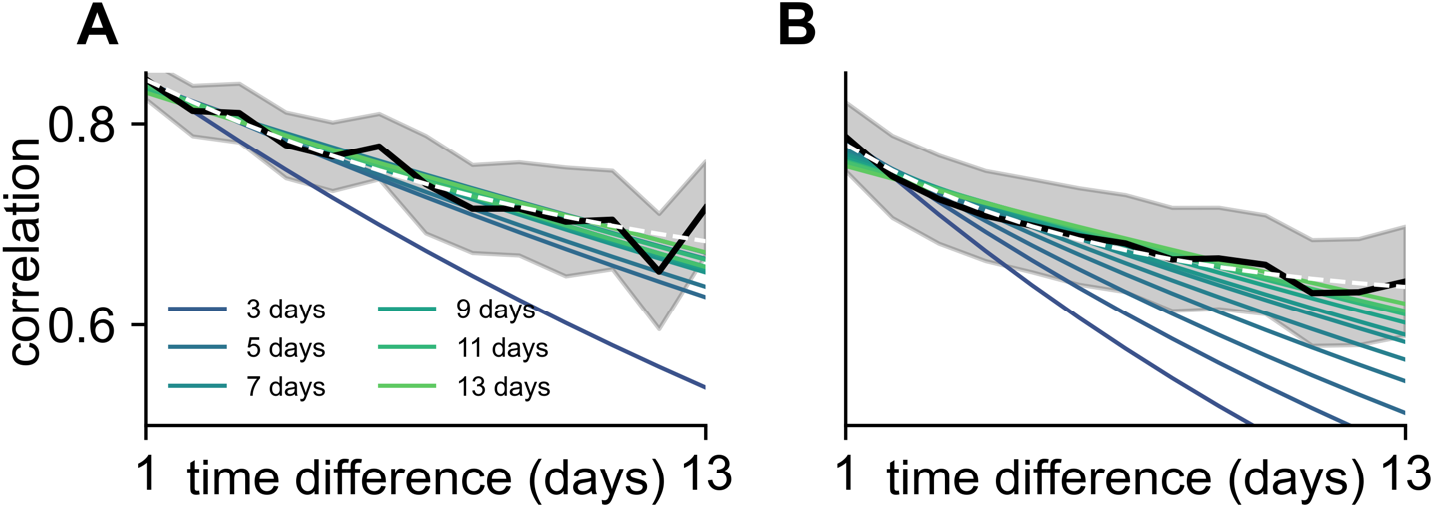
Exponential fits for different subsampled recording durations. We fitted exponential models to the average data across neurons recorded for at least 14 days during the lever-pressing task (Figure 4C), but considering only data up to and including increasing time differences (legend). As the subsampled ‘recording duration’ increases, so does the stability index learned in the exponential model for both neurons recorded in DLS (A) and MC (B). White dashed lines indicate exponential model fits with a learnable baseline, which better capture the decreasing rates of decay with increasing recording duration.

**Extended Data Figure 9:**
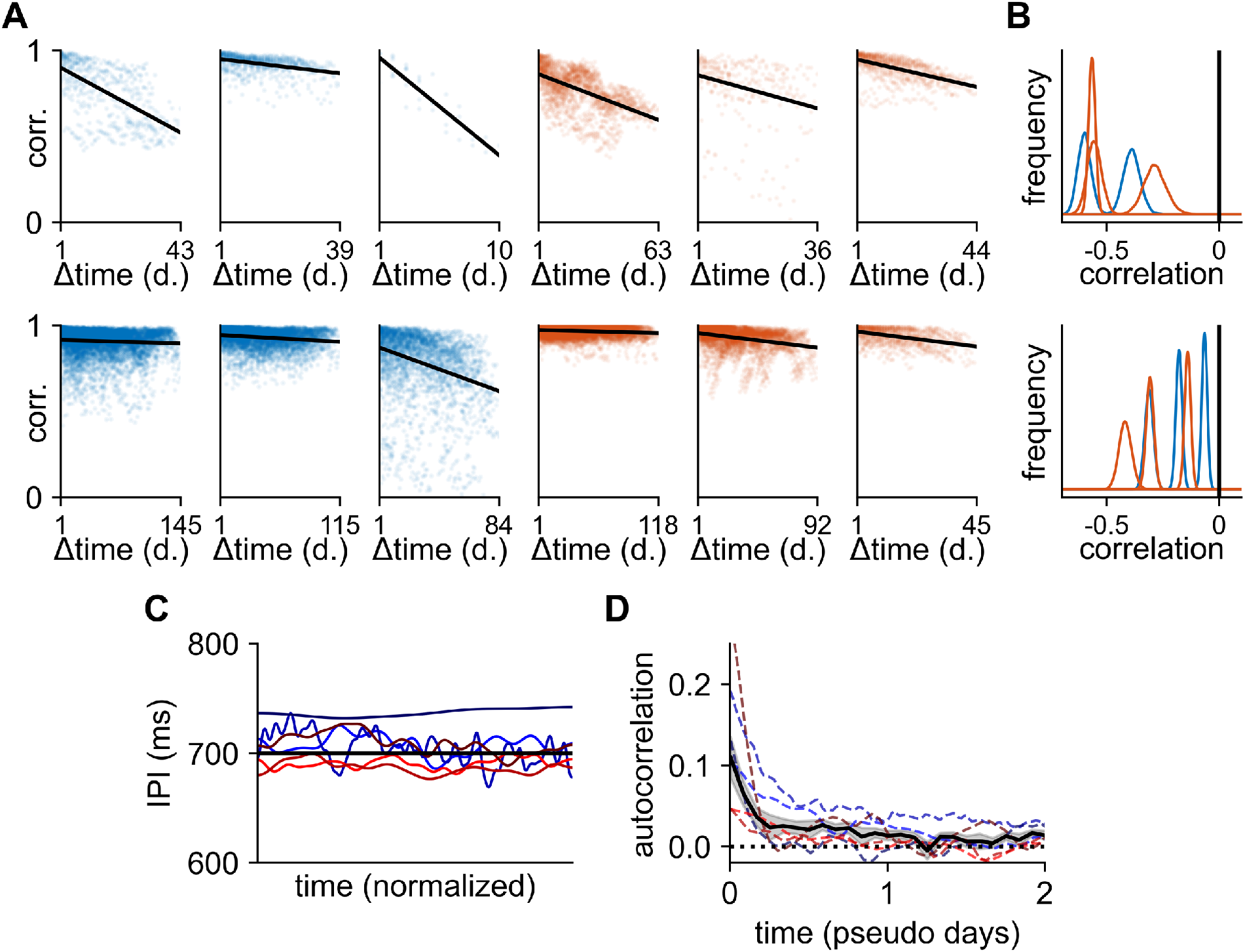
Behavioral drift and inter-press intervals. **(A)** Correlations between mean velocity profiles plotted against time difference for all pairs of days in each animal. Top row: lever-pressing task; bottom row: wet dog shakes. Blue indicates animals with recordings from DLS, red from MC. **(B)** Distribution of correlations between time difference and behavioral similarity across all animals, generated by a bootstrap analysis of the data in (A). All animals exhibit a significant negative correlation between behavioral similarity and time difference in both the lever-pressing task and wet dog shake behavior (p < 0.001; bootstrap test). **(C)** Inter-press interval (IPI) for each animal, convolved with a 200-trial Gaussian filter. Time is normalized from 0 to 1 for each animal (n = 9365 ± 6886 trials, mean ± std). Black horizontal line indicates 700 ms. **(D)** We computed the IPI autocorrelation as a function of trial number and normalized time by the average number of trials per day for each animal (colored lines). Black line and shading indicate mean and standard error across animals. Task performance is only correlated over short timescales of 0.5-1 days despite behavioral drift on timescales of weeks (c.f. panel A). This suggests that behavioral changes are predominantly along ‘task-null’ directions that do not affect performance.

**Extended Data Figure 10:**
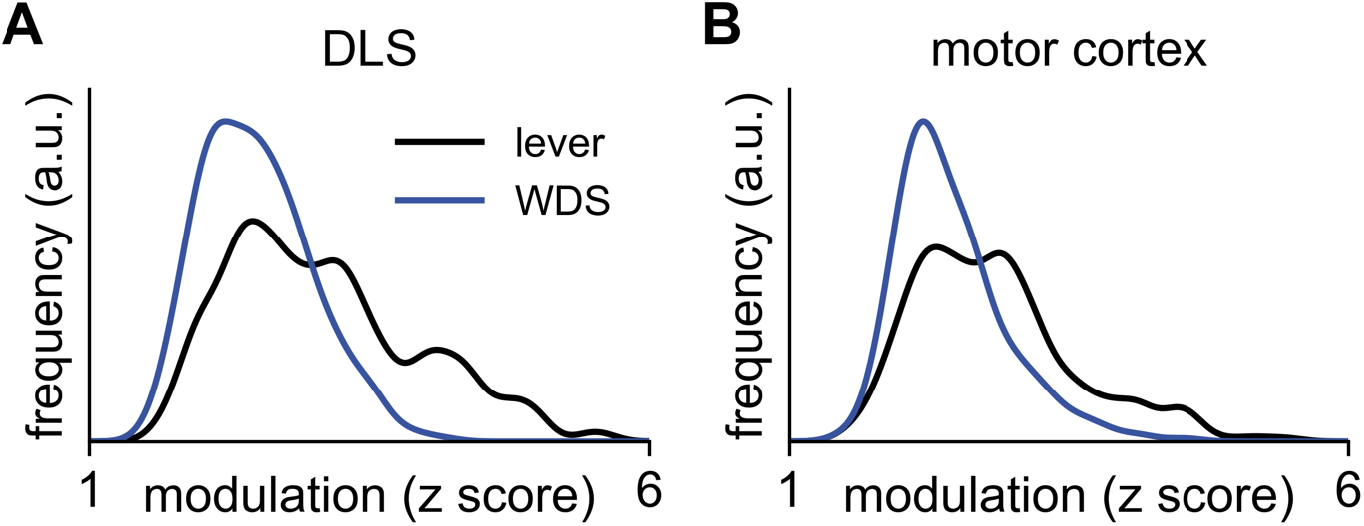
Task-modulation of neurons in the lever-pressing task and wet-dog shake behavior. **(A)** A PETH was computed across all trials for each neuron in 20 ms bins, and the time bin identified with the maximum deviation from the mean across all time bins. The corresponding z-score was computed, and the distribution of absolute values of these z-scores plotted across all DLS neurons for the lever-pressing task (black) and wet-dog shake behavior (blue). **(B)** As in (A), now for neurons recorded from MC.

## Notes

### Competing Interest Statement

The authors have declared no competing interest.

